# Oncolytic peptide NF27 effectively inhibits tumor growth and eradicates tumors in multiple cancer types

**DOI:** 10.1101/2025.08.17.668569

**Authors:** Natsuki Furukawa, Alex R. Chao, Wendy Yang, Akash Patil, Eric S. Christenson, Adam C. Mirando, Niranjan B. Pandey, Aleksander S. Popel

## Abstract

Oncolytic peptides are amphipathic peptides that specifically induce cell death in cancer cells by rupturing the cell membrane. Despite their therapeutic potential, few have advanced to clinical trials, and none have been approved for cancer treatment, highlighting the need for more potent and safe candidates. Moreover, the structure-activity relationship (SAR) of oncolytic peptides remains poorly understood. To address these challenges, we designed a series of peptides based on the previously reported oncolytic peptide CKS1 and evaluated their activity to induce cancer cell death. By comparing the structures and the activities of these peptides, we discovered novel insights in the SAR of oncolytic peptides. Among the peptides, we identified NF27 as the most potent peptide. NF27 showed broad cytotoxicity across multiple cancer types but displayed minimal toxicity against healthy cells and low hemolysis. Cell death induced by NF27 was immunogenic and promoted infiltration of immune cells in murine tumors. In murine tumor models, NF27 effectively suppressed tumor growth and achieved complete eradication in some cases, with no observable side effects. These findings highlight NF27 as a promising lead peptide for the development of safe and effective oncolytic therapies.

**One Sentence Summary:** We identified NF27, a novel oncolytic peptide that induces immunogenic cancer cell death and eradicates tumors with low toxicity via SAR study.

## INTRODUCTION

Cancer is responsible for nearly 10 million deaths annually worldwide (*1*). Despite recent advances in targeted therapy and immunotherapy, systemic chemotherapy remains the mainstay of treatment for many cancer patients (*2*). Systemic chemotherapy inhibits not only the proliferation of cancer cells but also healthy cells, particularly rapidly dividing cells such as those in the bone marrow, gastrointestinal tract, and hair follicles, resulting in a narrow therapeutic index (*3*). The damage to these healthy cells induces adverse side effects such as nausea, vomiting, diarrhea, loss of appetite, anemia, hair loss, fatigue, and immunosuppression. Because the kidney and liver are exposed to chemotherapy drugs during metabolism and elimination, nephrotoxicity and hepatotoxicity are also commonly seen during chemotherapy treatment. Furthermore, cancer cells can develop drug resistance via multiple mechanisms, including enhanced efflux of drugs due to the overexpression of transporters, upregulation of DNA repair mechanisms, inhibition of apoptosis, and the acquisition of detoxification mechanisms that neutralize the reactive intermediates (*4*). Given these limitations, there is an urgent need for safe and effective cancer treatments.

Oncolytic peptides, also known as anticancer peptides, are amphipathic, cationic peptides that induce cancer cell death by disrupting the cell membrane integrity (*5, 6*). These peptides interact with phospholipids in the cell membrane and undergo a conformational change to induce membrane pore formation. Their specificity for cancer cells arises from the differences in membrane composition between cancerous and healthy cells. Unlike healthy cell membranes, which are typically neutrally charged, cancer cell membranes are negatively charged. Chen et al. quantified the cell surface charge of commonly used cell lines using electrostatically charged nanoprobes and demonstrated that all 22 cancer cell lines tested exhibited a negative charge, whereas all 4 primary healthy cells showed neutral or slightly positive charges (*7*). Several factors contribute to the negative charge of cancer cell membranes, including the exposure of phosphatidylserine, a negatively charged phospholipid, to the outer cell membrane; the upregulation of anionic glycans such as sialic acids and glycosaminoglycans due to dysregulated glycosylation pathways; and the excessive production of lactate caused by altered cancer cell metabolism (*5, 7*). Since oncolytic peptides are positively charged, they preferentially bind to cancer cells over healthy cells. Membrane fluidity also plays a role in the sensitivity to oncolytic peptides. Compared to cancer cells, healthy cell membranes typically contain higher levels of cholesterol, which decreases membrane fluidity and reduces peptide absorption and insertion into the lipid bilayer. This contributes to the greater resistance of healthy cells to oncolytic peptides (*5, 8*).

Oncolytic peptides offer several therapeutic advantages. First, they selectively target cancer cells based on membrane properties rather than proliferation rates, reducing the risk of stem cell toxicity and associated side effects. Second, their mechanism of action does not rely on specific signaling pathways like proliferation or apoptosis. This minimizes the likelihood of drug resistance and enables oncolytic peptides to effectively target heterogeneous cancer cell populations, overcoming a major limitation of conventional therapies. Third, the short half-life of oncolytic peptides in serum reduces the risk of systemic toxicity. Lastly, oncolytic peptides promote anti-cancer immunity by inducing the release of neoantigens and damage-associated molecular patterns (DAMPs), enhancing immune system activation against tumors (*9*).

To date, four oncolytic peptides (CyPep-1, EP-100, LL-37, LTX-315) have been evaluated in clinical trials (*10*). EP-100 is a conjugate of a 10 amino acid natural agonist peptide of luteinizing hormone-releasing hormone (LHRH) and CLIP-71, an 18 amino acid lytic peptide. In the clinical trial of EP-100, the drug was administered intravenously for cancer types expressing LHRH, including breast and ovarian cancer (*11*). LL-37 was originally identified as an antimicrobial peptide (AMP) secreted by human granulocytes (*12*). It was later discovered that LL-37 can induce cell death in cancer cells (*13*). LTX-315 was designed based on a host defense peptide, lactoferricin (*14*). Szczepanski et al. created a library of 96 peptides derived from the active sites of known tumor suppressors and oncogenes, and identified several peptides with significant cytotoxic effects (*15*). Through structural optimization, they developed CyPep-1. In clinical trials, CyPep-1, LL-37, and LTX-315 were all used as intratumorally injected drugs.

The discovery of AMPs dates back to 1922 with the discovery of lysozyme by Alexander Fleming (*16*). Owing to decades of research, thousands of AMPs have been identified, and extensive structure-activity relationship (SAR) studies have provided a rich understanding of their biochemical properties (*17–19*). Recent high-throughput screening and machine learning approaches have further expanded this knowledge base (*20–22*). In contrast, oncolytic peptides have received comparatively little attention. While several AMPs have been shown to possess oncolytic activity, these findings have largely been framed as extensions of AMP research rather than focused efforts to develop cancer therapeutics. Importantly, most of these studies have not addressed the central challenge in oncolytic peptide development: to selectively kill cancer cells while sparing healthy mammalian tissues. This challenge differs fundamentally from that of AMPs, where the goal is to target bacteria, which are organisms that are evolutionarily and structurally distinct from mammalian cells.

Despite the therapeutic potential, no oncolytic peptide has yet received regulatory approval for cancer treatment. A major limitation is the scarcity of SAR studies, which are crucial for guiding the rational design of safer and more effective oncolytic peptides. To address this gap, we designed a series of oncolytic peptides based on CKS1, a previously reported peptide with anti- angiogenic and oncolytic properties (*23*). By comparing the predicted structures of these peptides, we identified key structural features that influence oncolytic efficacy. This approach led to the identification of NF27, a peptide that exhibits potent and selective oncolytic activity. We subsequently investigated mechanism of action of NF27 through a series of cellular assays and evaluated the therapeutic potential of NF27 in preclinical cancer models.

## RESULTS

### Design and assessment of CKS1-derived oncolytic peptides

To investigate the SAR of oncolytic peptides, we designed a series of peptides derived from CKS1. CKS1 consists of an α-helical region and an unstructured region separated by proline residues. We previously reported that the α-helical region is indispensable for the oncolytic activity of CKS1 (*23*). Therefore, we first attempted to optimize the structure of the α-helical region. To enhance the positive charge necessary for targeting cancer cell surfaces, we substituted glutamic acid (E19) with lysine (NF04). We further designed peptides with increased positive charge and hydrophobicity, respectively, by replacing neutrally charged polar amino acids with lysine and isoleucine. We also replaced methionine (M21) with other amino acids since the oxidative nature of sulfur decreases the stability of the peptides (NF05 – NF08). A previous SAR study of AMPs reported that amino acids with large aromatic side chains, such as tryptophan and phenylalanine, reduce selectivity and promote hemolysis (*21*). The same study also found that removing cysteine decreased hemolytic activity. Based on these findings, we intentionally avoided using these amino acids in our peptide design. CKS1 contains two prolines (P9 and P12) in its sequence, which disrupts the helical structure and causes kinks at those locations. To test the influence of these prolines on the oncolytic activity, we designed peptides that lack P12 (NF09 – NF13) and peptides that lack both proline residues (NF20 – NF22). These peptides had longer helical regions compared to CKS1. In particular, NF20 – NF22 became a single α-helix with no other secondary structure as illustrated by NF21 (**Fig. S1**). The helical wheel projections of the peptides are listed in **Fig. S2**. We further designed peptides with mutations in the non-helical region (NF14 – NF19) to test whether the characteristics of the non-helical region affect the potency of the peptide. Finally, we replaced the lysine residues with arginine residues to determine whether there is a difference between lysine and arginine (NF23 – NF26). The sequence alignment of the peptides is shown in **Fig. S3**.

We tested the oncolytic activity of each peptide by treating 4T1 murine triple-negative breast cancer cells with six different concentrations of the peptide and measuring cell death via lactate dehydrogenase (LDH) assay (**Fig. 1A**). The half-maximal effective concentration (EC_50_) values were determined based on dose-response curves fitted to the experimental data. NF11 was excluded from the study due to its poor solubility, as it formed a hard gel in water, making it difficult to assess activity. While NF22 showed no detectable oncolytic activity in the initial screen, a subsequent test using a higher-purity lot confirmed activity, with an EC_50_ value of 78.9 µM, the weakest among the peptides tested (**Fig. S4**). The physical properties of the peptides, estimated using the modlAMP package (*24*), are summarized in **Data S1**.

**Fig. 1.**
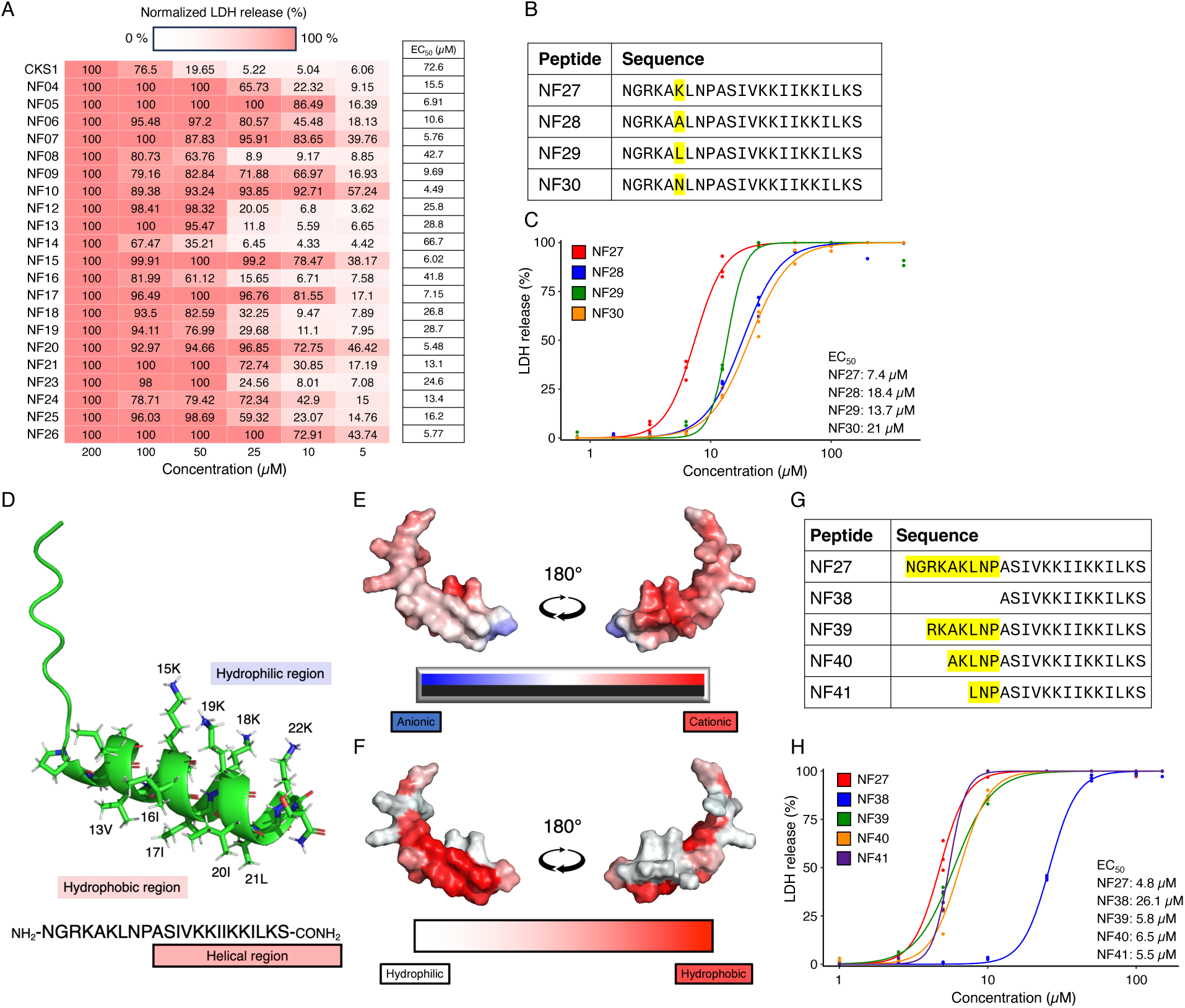
Oncolytic activity of the CKS1-derived peptides. (A) The LDH release from 4T1 cells treated with CKS1-derived peptides. LDH release was normalized to the maximum release observed at the highest peptide concentration. The EC_50_ values were determined by fitting dose-response curves to the experimental data. Representative of N = 2. (B) Sequence alignment of NF27-derived peptides. (C) Dose-response curves and corresponding EC₅₀ values of NF27- derived peptides. Representative of N = 3. Each dot represents a technical replicate. (D) Predicted structure of NF27. (E) The visualization of the electrostatic potential of NF27. (F) The visualization of the hydrophilic and hydrophobic regions of NF27. (G) Sequence alignment of NF27 and its truncated peptides. (H) Dose-response curves and EC_50_ values of NF27 and its truncated peptides. Each data point represents a technical replicate.

By comparing the structures and the activities of each peptide, we gained several insights into their SAR. First, positive charge is essential for the oncolytic activity; however, increasing positive charge does not necessarily enhance potency. In our previous study on CKS1, we demonstrated that removing positive charge from the helical region by replacing three lysine residues with glutamic acid residues completely abolished oncolytic activity. In the present study, we found that replacing E19 with a lysine residue increased oncolytic activity (**Fig. S5A**). However, after incorporating the results of all peptides, our correlation analysis revealed that charge density did not directly correlate with oncolytic activity (**Fig. S5B**).

Second, higher hydrophobicity correlated with greater oncolytic activity. For example, NF07, which has a larger hydrophobic region compared to NF08, was more potent than NF08 (**Fig. S6A**). Likewise, NF21, with hydrophobic residues at positions 6 and 7, exhibited markedly higher activity than NF22, which contains hydrophilic residues at the same positions (**Fig. S6B**).

We observed a negative correlation between the EC_50_ values and the aliphatic index, which is a measure of the relative volume occupied by the aliphatic side changes of hydrophobic amino acids (**Fig. S6C**). Similarly, EC_50_ values negatively correlated with the proportion of hydrophobic amino acids in the peptide (i.e. the relative frequency of alanine, cysteine, phenylalanine, isoleucine, leucine, methionine, and valine) (**Fig. S6D**).

Third, the role of proline residues and the length of the helical region appears to be more nuanced and context dependent. In several cases, removing P12 and increasing the length of the helical region enhanced oncolytic activity (**Fig. S7A, B**). Eliminating both P12 and P9 to generate a fully helical structure also improved potency (**Fig. S7C, D**). However, removing the P9 resulted in a decrease of activity in the case of NF22. NF19, which retains P9 and thus contains a structural kink, exhibited significantly greater oncolytic activity than NF22, which lacks proline and forms a helical conformation (**Fig. S7E**).

We next focused on optimizing NF10, the most potent peptide among the CKS1-derived peptides. As NF10 contains cysteine, a residue that can compromise stability and specificity, we attempted to replace it with other amino acids. We designed four peptides (NF27 – NF30), with lysine, alanine, leucine, or asparagine in place of the cysteine residue. Among these peptides, NF27 exhibited the highest potency (**Fig. 1B, C**). Structurally, NF27 retains key features of the original CKS1 peptide, including an unstructured N-terminal region and an amphipathic α-helix at the C-terminus. Removal of a proline residue resulted in a longer helical region. While CKS1’s hydrophilic face includes glutamic acid and asparagine residues, the hydrophilic region of NF27 is composed exclusively of lysine residues, contributing to a higher net positive charge and a more distinct segregation between hydrophilic and hydrophobic domains (**Fig. 1D – F**). To evaluate whether the hydrophobic amino acids could be further optimized, we substituted them with phenylalanine, a commonly used hydrophobic residue in oncolytic peptides and AMPs.

However, this substitution did not enhance potency compared to NF27 (**Fig. S8A, B**). Based on this finding, we chose to retain isoleucine and leucine residues to construct the hydrophobic region.

To evaluate the functional role of the unstructured N-terminal region, we generated truncated peptides (NF38 – NF41) and assessed their oncolytic activity. While NF39 – NF41 showed no significant difference in potency compared to NF27, NF38, which completely lacks the N-terminal region and adopts a singular helical structure, exhibited significantly reduced activity (**Fig. 1G, H**).

Previously, we reported that the amphipathic structure of the helical region is critical for the oncolytic activity of CKS1, and that disrupting this amphipathic helix diminishes activity. To determine whether this structural requirement also applies to NF27, we designed two variants: NF42, in which a proline residue was inserted into the helical region to disrupt the secondary structure, and NF43, in which two hydrophobic isoleucine residues (I16 and I17) were replaced with hydrophilic lysine residues (K16 and K17). NF42 exhibited minimal oncolytic activity at concentrations above 500 µM, whereas NF43 exhibited no detectable activity even at 1000 µM (**Fig. S9A – D**). These findings confirm that the amphipathic helical structure of NF27 is critical for its function, with the hydrophobic isoleucine residues playing a particularly important role.

Finally, we evaluated the impact of C-terminal amidation on NF27’s activity. C-terminal amidation is a common modification in natural AMPs and is believed to enhance activity through several mechanisms, including increased net positive charge, stabilization of α-helix structure via additional hydrogen bonding, and protection from carboxypeptidase-mediated degradation (*25, 26*). To assess the role of C-terminal amidation in NF27, we synthesized a non-amidated version and evaluated its oncolytic activity against 4T1 cells. While the non-amidated peptide retained substantial potency, the EC_50_ value was 19.0 µM, higher than that of the amidated form, indicating that amidation enhances NF27’s potency (**Fig. S9E**).

### Oncolytic activity of NF27

To evaluate the efficacy and specificity of NF27 to induce cancer cell death, we tested NF27 against multiple cell lines and quantified the cell death induced by the peptide. In both murine and human cell lines, NF27 induced cell death in every cancer cell line tested (**Fig. 2A, 2B**). The EC_50_ values of NF27 against non-cancerous cells, namely 3T3 fibroblasts, human umbilical vein endothelial cells (HUVECs), and normal human lung fibroblasts (NHLFs), were significantly higher compared to those against cancerous cells. We further validated the cytotoxic activity of NF27 by alamarBlue cell viability assay (**Fig. S10**).

**Fig. 2.**
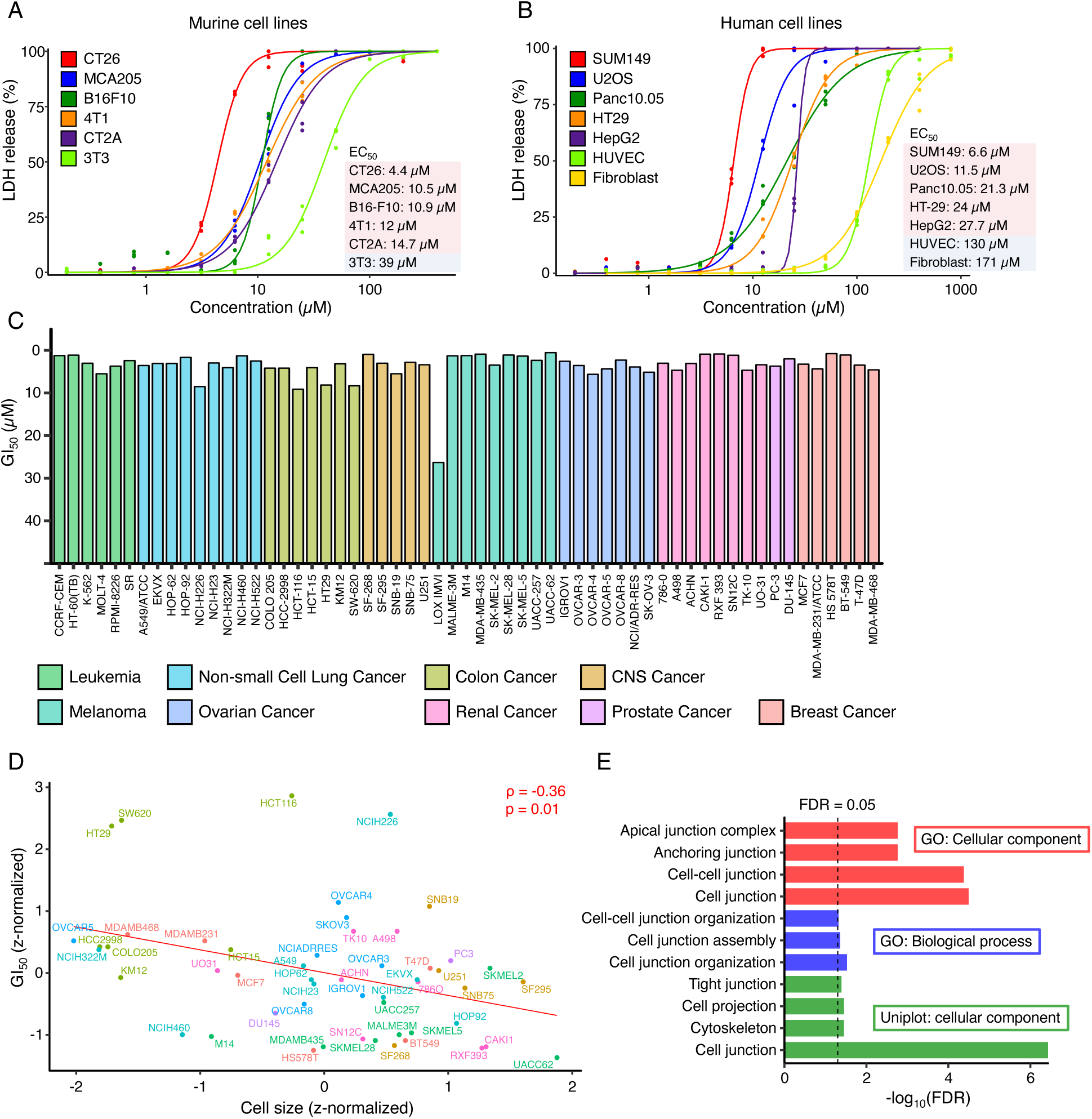
NF27 exhibits broad-spectrum cytotoxicity against cancer cell lines. (A) Dose- response curves and EC₅₀ values of NF27 in murine cancer cell lines. Representative of N = 3. Each data point represents a technical replicate. (B) Dose-response curves and EC₅₀ values of NF27 in human cancer cell lines. Representative of N = 3. Each data point represents a technical replicate. (C) GI₅₀ values for NF27 across 59 human cancer cell lines from the NCI-60 panel. (D) Correlation between GI₅₀ values and average cancer cell size. (E) Enriched gene ontology (GO) terms among genes showing significant positive correlation with NF27 GI₅₀ values.

To evaluate the oncolytic activity of NF27 against a broader range of cancer cells, we utilized the NCI-60 human tumor cell lines screen. One cell line was removed from the screen due to a quality control issue. Among the remaining 59 human cancer cell lines tested, the GI_50_ values (the concentration needed to inhibit 50% of cell growth) were consistently below 10 µM (**Fig. 2C, Data S2**). There was one exception, human melanoma cell line LOX IMVI, which had a GI_50_ value of 26.3 µM.

To determine why certain cell types respond better than others to NF27, we assessed the correlation between the sensitivity to NF27 and various cellular features. Specifically, we collected morphological data and omics data using the NCI-60 cancer cell lines and calculated the correlation metrics. We observed a negative correlation between the cell size (*27*) and the GI_50_ values, indicating that larger cells tend to be more sensitive to NF27 (**Fig. 2D**). Using transcriptomics data (*28*), we extracted genes with expression levels that were significantly correlated with the GI_50_ values at a false discovery rate of 5%. We identified 110 genes with expression levels that positively correlated with the GI_50_ values and 20 genes with expression levels that negatively correlated with the GI_50_ values (**Data S3**). We conducted a gene enrichment analysis to identify the features that correlate with sensitivity to NF27. We discovered that genes related to cell junctions were highly expressed in cell lines that are relatively resistance to NF27 (**Fig. 2E, Fig. S11, Fig. S12**). We also observed a correlation between GI_50_ values and the expression of epithelial cell adhesion molecule (EpCAM) at the protein level (**Fig. S13**).

### Mechanism of cell death caused by NF27

To study the mechanism of cell death induced by NF27, we stained 4T1 cells with Calcein-AM and observed the morphology after NF27 treatment. NF27 caused pore formation in the cell membrane, leading to the leakage of the cytosol (**Movie S1**). In some instances, pore formation resulted in the influx of surrounding fluid, causing the formation of blebs. Eventually, the blebs burst, leading to cytosol leakage and cell death (**Fig. 3A, Movie S2**). To quantify the speed of NF27-induced membrane rupture, we measured the fraction of Calcein^+^ cells over time. The fraction of Calcein^+^ cells began to rapidly decrease at approximately 10 minutes after NF27 treatment and reached 0% at approximately 20 minutes after treatment (**Fig. 3B**). These data demonstrate that NF27 rapidly induces membrane damage upon treatment.

**Fig. 3.**
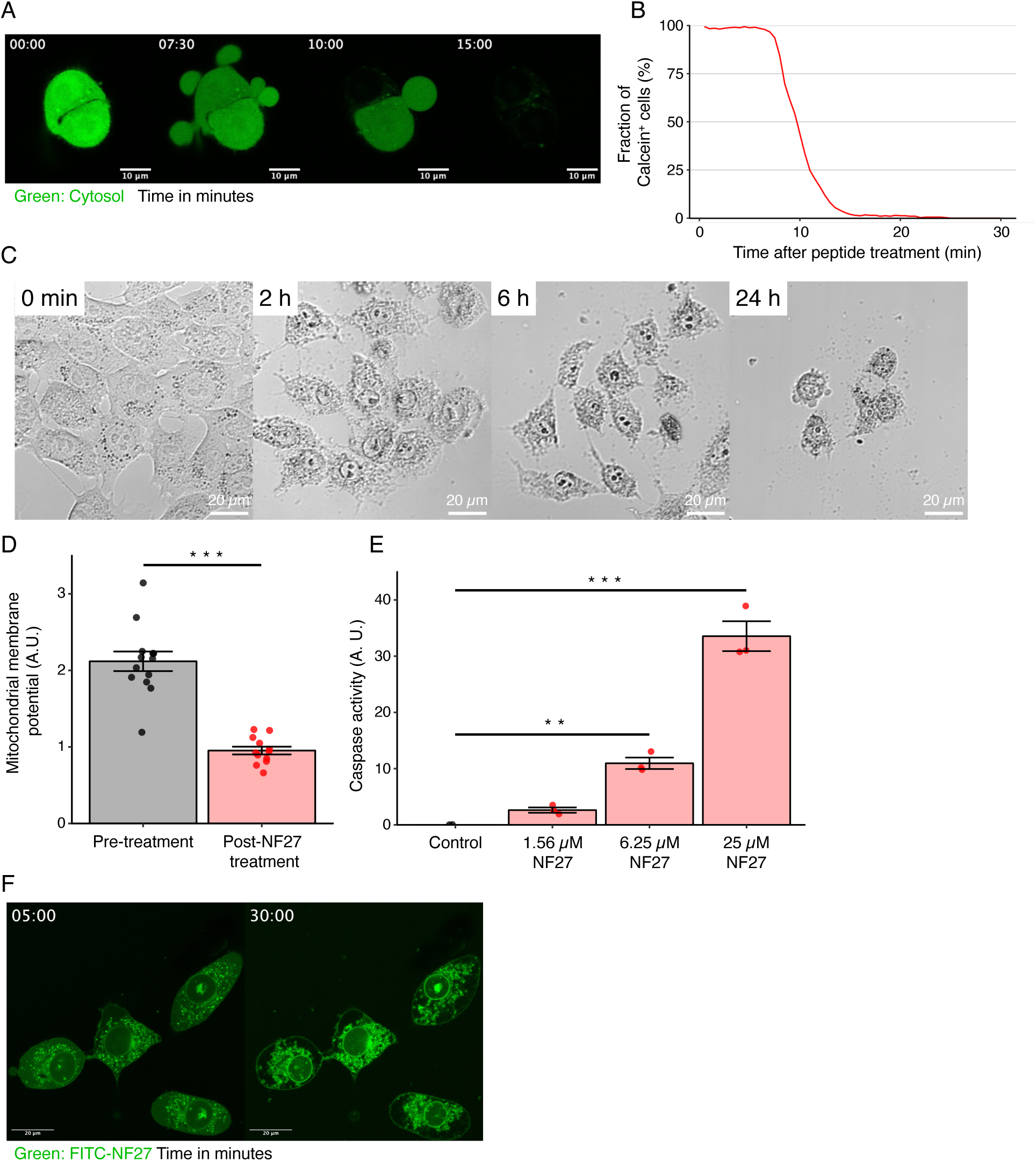
Mechanism of NF27-induced cell death. (A) Live-cell imaging of Calcein-AM-stained 4T1 cells following 20 µM NF27 treatment. (B) Quantification of Calcein^+^ 4T1 cells post- treatment. Representative of N = 3. (C) Brightfield images showing morphological changes in 4T1 cells after 20 µM NF27 exposure. (D) Analysis of mitochondrial membrane potential in JC- 1-stained 4T1 cells treated with 20 µM NF27. The y-axis represents the ratio of red fluorescence to green fluorescence. N = 3; data are presented as mean ± SEM. Each data point represents the red/green ratio of an individual cell. Two-tailed Welch’s t-test was performed to determine the significance (***p < 0.005). (E) Activation of caspase-3 after 6 hours of NF27 treatment. N = 3; data are presented as mean ± SEM. Each data point represents the mean of triplicate measurements from each experiment. Dunnet’s test was performed to determine the significance (**p < 0.01, ***p < 0.005). (F) Fluorescence microscopy image showing the intracellular localization of FITC-labeled NF27 in 4T1 cells. 4T1 cells were treated with 10 µM FITC-NF27.

We observed two distinct fates in cells treated with NF27. Most cells detached from the culture plate shortly after exposure, while a subset remained attached even after prolonged exposure. These cells exhibited apoptotic morphological features such as cell shrinkage, chromatin condensation, and the collapse of the nucleus (**Fig. 3C**). Based on these observations, we hypothesized that NF27 damages the cell membrane while simultaneously activating the apoptotic pathway. When membrane damage is severe, necrotic cell death likely predominates, resulting in rapid detachment. If cells survive the initial membrane damage, they ultimately undergo apoptotic cell death following entry of the peptide into the cell. To test the involvement of the apoptotic pathway, we measured the mitochondrial membrane potential and caspase activation following NF27 treatment. The mitochondrial membrane potential was rapidly lost after NF27 treatment (**Fig. 3D, Movie S3**). Dose-dependent caspase-3 activation was observed six hours post-treatment (**Fig. 3E**), indicating the activation of the apoptotic pathway.

To further explore the intracellular localization of NF27, we treated 4T1 cells with fluorescein isothiocyanate (FITC)-labeled NF27. After entering the cell, the peptide localized to multiple structures, including the plasma membrane, nuclear envelope, and nucleus (**Fig. 3F, Movie S4**). Initially distributed throughout the cytosol, FITC-NF27 later disappeared from this compartment, which may be a result of either accumulation in intracellular organelles or cytosolic leakage due to membrane disruption. To determine potential organelle specificity, we examined colocalization of FITC-NF27 with markers for mitochondria, the Golgi apparatus, and the endoplasmic reticulum (**Fig. S14A – C**). While partial colocalization was observed with all three markers, NF27 did not preferentially accumulate in any specific organelle (**Fig. S14D**).

This localization pattern differs from that of other oncolytic peptides, such as LTX-315 and LTX-401, which have been reported to selectively target mitochondria (*29*) and the Golgi apparatus (*30*), respectively.

### Immunogenicity of cell death caused by NF27

Oncolytic peptides, including CKS1, induce the release of immunogenic molecules (*23*). To determine whether NF27 has a similar effect, we quantified the amount of ATP release from NF27-treated cells. ATP functions as a damage-associated molecular pattern (DAMP), an indicator of immunogenic cell death (*31*). NF27 significantly induced ATP release from both 4T1 and CT26 cells (**Fig. 4A, 4B**). To assess whether immunogenic molecules released from NF27-treated cancer cells activate immune cells, we exposed DC2.4 murine dendritic cells to conditioned media from NF27-treated cancer cells. Treatment with conditioned media from both 4T1 and CT26 cells increased the release of TNF-α and IL-6, indicating dendritic cell activation (**Fig 4C, 4D**).

**Fig. 4.**
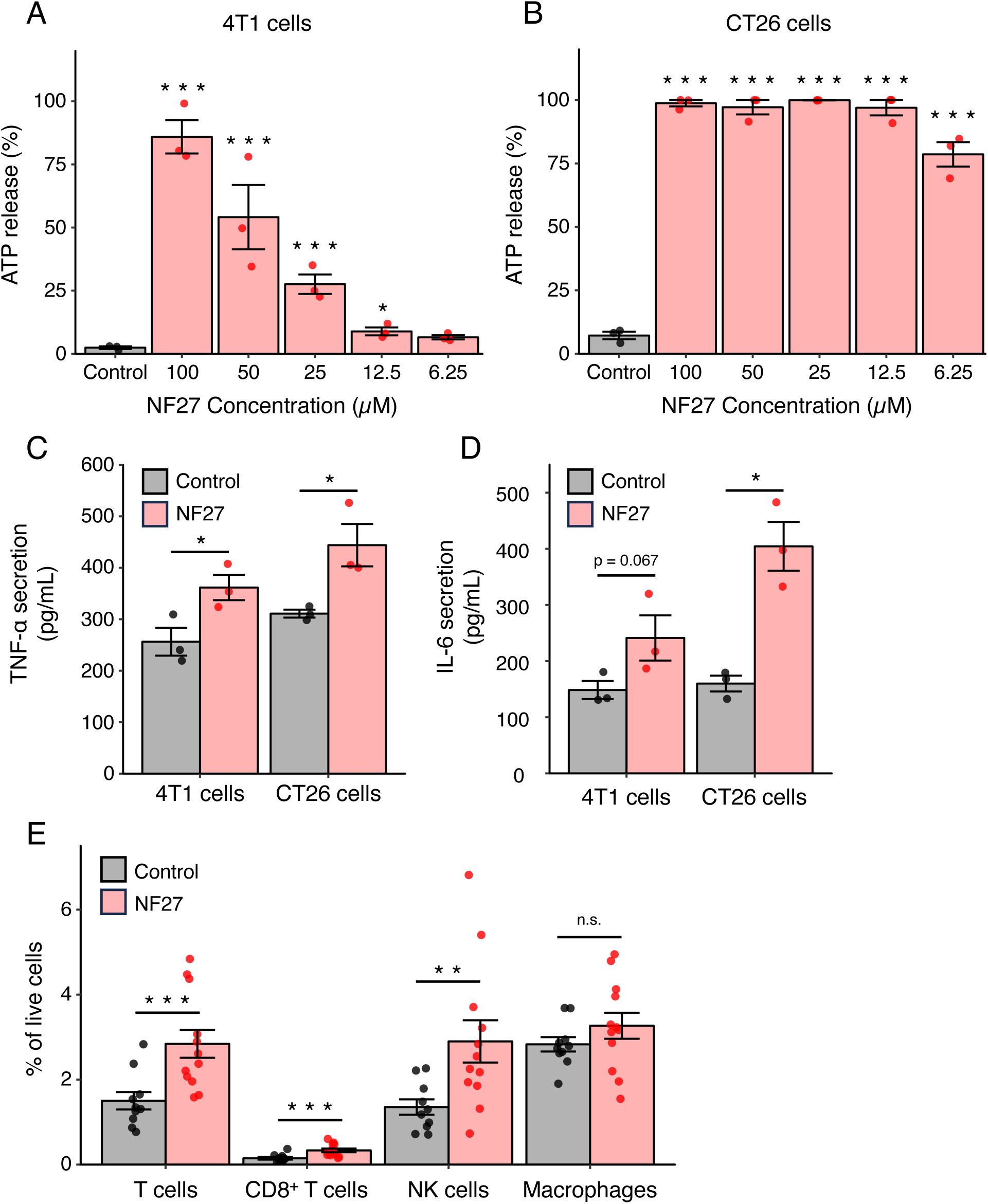
NF27 induces immunogenic cell death. (A, B) ATP release from NF27-treated 4T1 and CT26 cells. ATP release was normalized to the maximum release observed in cells treated with lysis buffer. N = 3: data are presented as mean ± SEM. Each data point represents the mean of triplicate measurements from each experiment. Dunnet’s test was performed to determine the significance (*p < 0.5, ***p < 0.005). (C) TNF-α secretion by DC2.4 dendritic cells following treatment with conditioned media from NF27-treated cancer cells. N = 3 per group; data are presented as mean ± SEM. Each data point represents the mean of triplicate measurements from each experiment. Two-tailed Welch’s t-test was performed to determine significance (*p < 0.05). (D) IL-6 secretion by DC2.4 dendritic cells following treatment with conditioned media from NF27-treated cancer cells. N = 3 per group; data are presented as mean ± SEM. Each data point represents the mean of triplicate measurements from each experiment. Two-tailed Welch’s t-test was performed to determine significance (*p < 0.05). (E) Flow cytometry analysis of CT26 tumors treated with vehicle or NF27, showing the frequency of T cells (CD45⁺CD3⁺), CD8⁺ T cells (CD45⁺CD3⁺CD8⁺), NK cells (CD45⁺CD49b⁺CD3⁻), and macrophages (CD45⁺F4/80⁺CD11b⁺) among live cells. N = 10 (control), N = 12 (NF27); data are presented as mean ± SEM. Each data point represents the data from an individual mouse. Two-tailed Welch’s t-test was performed to determine significance (**p < 0.01, ***p < 0.005, n.s., not significant).

To evaluate the immunogenic effect of NF27 in vivo, we intratumorally injected vehicle or NF27 into CT26 tumors established in mice. Five days later, the tumors were excised and analyzed by flow cytometry (**Fig. 4E, Fig. S15**). NF27 treatment increased infiltration of T cells (CD45^+^CD3^+^), CD8^+^ T cells (CD45^+^CD3^+^CD8^+^), and NK cells (CD45^+^CD49b^+^CD3^-^). No significant change was observed in the infiltration of macrophages (CD45^+^ F4/80^+^CD11b^+^). These findings suggest that NF27 activates both innate and adaptive immune responses via the release of immunogenic molecules.

### NF27 inhibits tumor growth and eradicates tumors *in vivo*

To evaluate the therapeutic efficacy of NF27, we intratumorally injected 40 mg/kg NF27 in established tumors every other day and monitored tumor growth. NF27 significantly inhibited tumor growth across all tested models, including the CT26 murine colorectal model, the HT-29 human colorectal model, the B16-F10 murine melanoma model, and the 4T1 murine triple-negative breast cancer model (**Fig. 5A – D**). NF27 was particularly effective in CT26 models and HT29 models, where complete tumor eradication was achieved in 80% and 60% of treated mice, respectively (**Fig. S16A, B**). **Fig. S16C** shows a representative image of HT-29 tumor treated with NF27, which reveals pronounced tumor necrosis after a single dose of NF27.

**Fig. 5.**
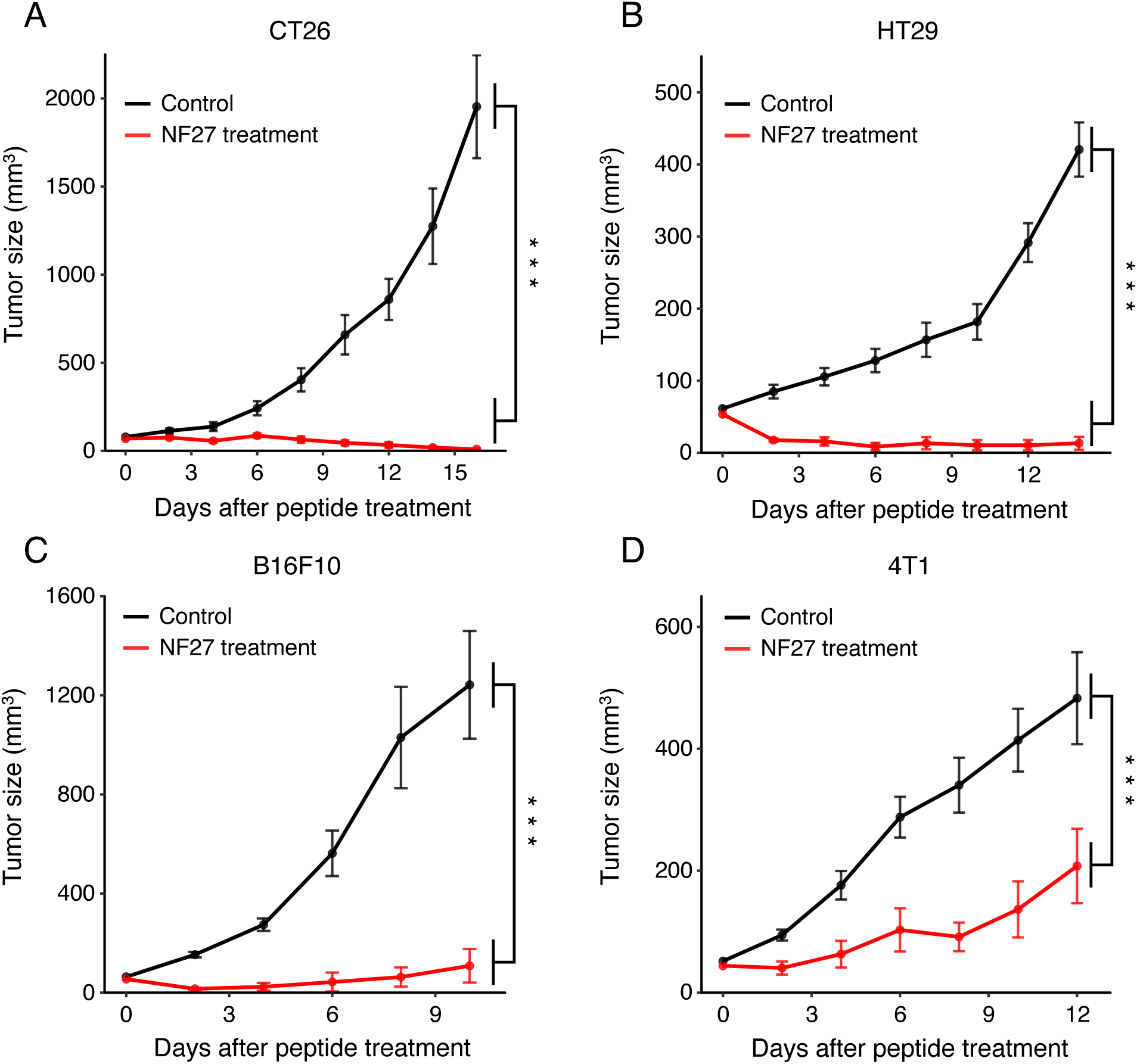
NF27 significantly inhibits tumor growth and eradicates tumors. (A–D) Tumor growth curves for CT26 (control, N = 10; NF27-treated, N = 10), HT-29 (control, N = 12; NF27- treated, N = 11), B16-F10 (control, N = 10; NF27-treated, N = 9), and 4T1 (control, N = 8; NF27-treated, N = 8) following NF27 treatment. Mice received intratumoral injections of 40 mg/kg NF27 every 2 days starting on day 0. Data are shown as mean ± SEM. Statistical significance was determined using two-way ANOVA (***p < 0.005).

### Translational potential of NF27

To assess the potential of NF27 as a cancer drug, we evaluated the stability of the peptide. We first calculated the instability index, which assigns a score based on the dipeptide patterns frequently observed in unstable proteins (*32*). Proteins with an instability index above 40 are considered to be unstable. NF27 had an instability index of -4.9 (**Data S1**), indicating high stability. To experimentally assess the stability of NF27, we solubilized NF27 in water at a concentration of 40 mM and incubated at either room temperature, 4°C, or -80°C for 30 days. After incubation, we evaluated the oncolytic activity using LDH assay. Even at room temperature, NF27 retained its activity after 30 days (**Fig. 6A**). We next analyzed the half-life of NF27 in serum using mass spectrometry. In murine serum, NF27 had a half-life of approximately 30 minutes when incubated at 37°C (**Fig. 6B**). These data demonstrate the physicochemical stability of NF27, which is crucial for storage and integration with drug delivery technologies. Additionally, the short half-life of NF27 in serum suggests a low risk of systemic toxicity, as it is rapidly degraded upon entering the bloodstream. Lastly, we evaluated the hemolytic activity of NF27, a common challenge for oncolytic peptides that limit their translational potential (*33*). We compared NF27 with melittin, which is an oncolytic peptide known to have hemolytic activity (*34*). Melittin demonstrated toxic hemolysis, defined as the lysis of more than 10% of red blood cells, at a concentration of 2 µM. In contrast, NF27 did not exhibit toxic hemolysis even at a concentration of 1000 µM (**Fig. 6C**).

**Fig. 6.**
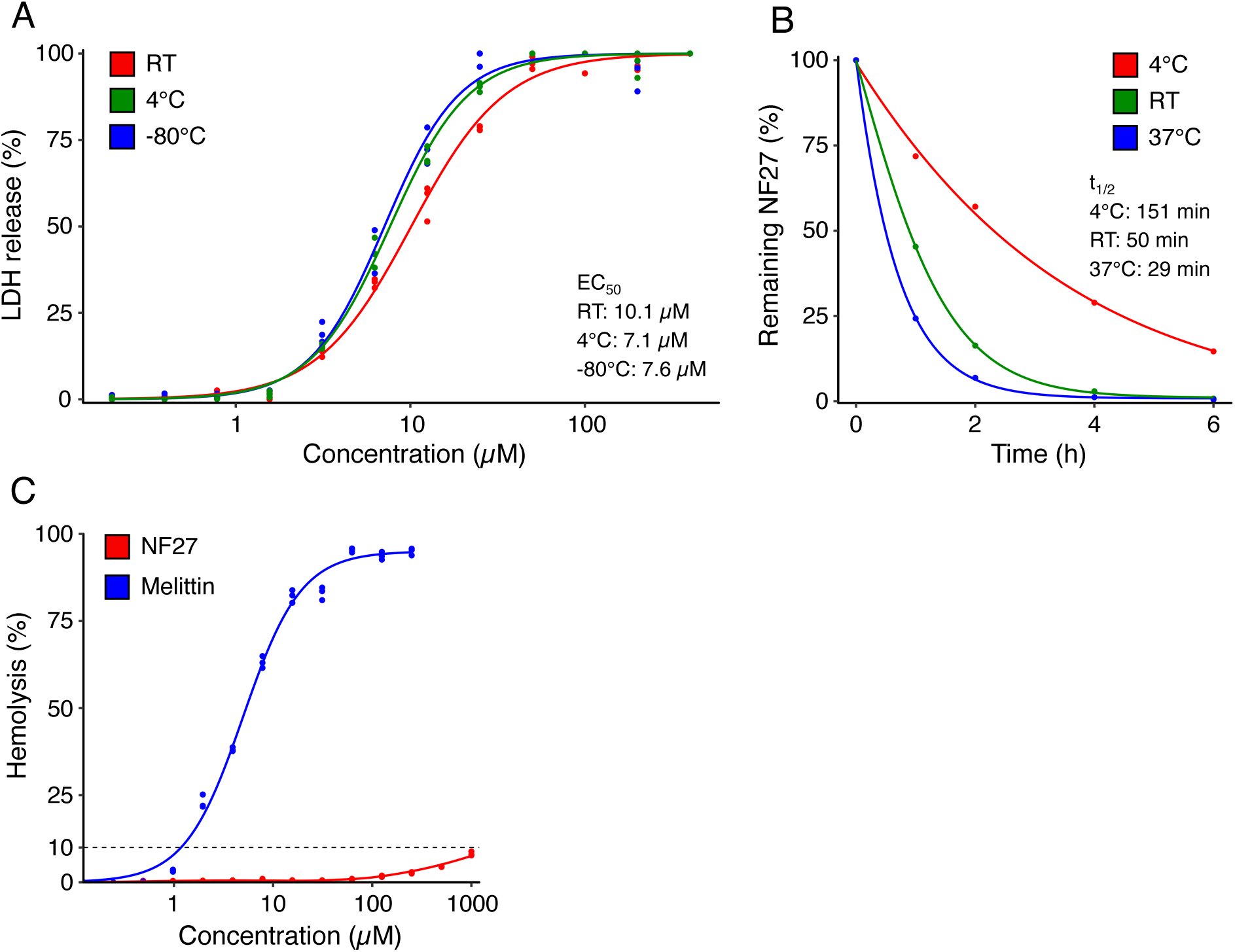
NF27 exhibits favorable pharmacological properties. (A) Stability of NF27 after 30- day incubation. NF27 was dissolved in water and stored at the indicated temperatures. Oncolytic activity against 4T1 cells was assessed using an LDH assay. Representative of N = 3. Each data point represents a technical replicate. (B) Pharmacokinetics of NF27 in murine serum. NF27 was incubated in murine serum at the indicated temperatures, and remaining peptide levels were quantified by mass spectrometry. (C) Hemolytic activity of melittin and NF27. Red blood cells were incubated with melittin or NF27, and the extent of hemolysis was quantified. Representative of N = 3. Each data point represents a technical replicate.

## DISCUSSION

In summary, we designed a series of peptides derived from CKS1, analyzed their oncolytic effects, and identified key structural features important for oncolytic activity. While previous SAR studies have highlighted general characteristics such as hydrophobicity and positive charge, few have explored the three-dimensional structures of oncolytic peptides or assessed their SAR at the single amino acid level. Consistent with previous findings (*35*), we confirmed that increased hydrophobicity positively correlates with oncolytic activity. The relationship between positive charge and the oncolytic activity was more nuanced. We demonstrated that replacing negatively charged amino acids with positively charged amino acids significantly improved the oncolytic activity of CKS1-derived peptides, whereas removing positive charge diminished oncolytic activity (*23*). However, we did not observe a direct correlation between increased positive charge and enhanced potency. These findings suggest that while a certain level of positive charge is required for activity, increasing charge does not necessarily enhance efficacy.

The role of proline-induced kinks in α-helical peptides remains controversial, with some studies suggesting that the kink enhances pore-forming activity, while others report a reduction in membrane-disruptive function (*36*). In our SAR study, we also observed mixed outcomes: depending on the peptide, removal of proline residues led to either an enhancement or a reduction in oncolytic activity. A detailed computational study by Tuerkova et al. revealed that kinks can destabilize barrel-stave pores and thereby reducing the activity of oncolytic peptides that act through this pore-forming mechanism. On the other hand, added flexibility introduced by a kink enhanced the oncolytic activity of peptides that form toroidal pores (*36*). We hypothesize that the physicochemical properties and structure of each peptide, which influence the pore-forming mechanism, likely determine the functional impact of proline-induced kinks.

In the case of NF22, the hydrophilic region and the hydrophobic region lie on the same side of the helix. Among the CKS1-derived peptides, NF22 exhibited the lowest oncolytic activity, potentially due to repulsive interactions between its hydrophilic residues and the lipid membrane. NF19, which includes a proline-induced kink that spatially separates the hydrophilic and hydrophobic regions, showed significantly higher activity. This suggests that such spatial segregation may reduce repulsive forces and enhance membrane interaction. NF21, which has increased hydrophobicity in the hydrophilic region, shows significantly greater activity than NF22. These observations suggest that the spatial arrangement of hydrophilic and hydrophobic residues and the resulting repulsive interactions, may be critical for oncolytic activity (**Fig. S17A**). NF27 features a highly hydrophobic face on one side of the helix, a highly hydrophilic face on the opposite side, and a proline residue that separates a highly hydrophilic nonhelical region (**Fig. S17B**). Our data demonstrate that NF27 exhibits higher oncolytic activity than NF38, which consists of only the helical region of NF27. This may be attributed to the reduced flexibility of NF38 or an active role of the unstructured hydrophilic region in facilitating membrane pore formation. Further studies are necessary to validate these hypotheses and to fully elucidate the SAR.

The discovery of inactive peptides NF42 and NF43 was particularly intriguing. These peptides differ from their oncolytic counterparts by only one or two amino acids and share similar overall physicochemical properties. Notably, NF27, NF42, and NF43 were all predicted to possess oncolytic activity by every machine learning-based predictor tested, including mACPpred 2.0 (*37*), ACPred-BMF (*38*), AntiCP 2.0 (*39*), and MLACP 2.0 (*40*). These findings highlight the limitations of sequence-based predictions and underscore the importance of incorporating three-dimensional structural features when evaluating peptide activity.

During the SAR study, we identified NF27 as a potent oncolytic peptide with high efficacy against cancer cells and high specificity. NF27 induced cell death in all cancer types tested, demonstrating pan-cancer activity. In addition to its direct cytotoxic effects, NF27 activated anti-cancer immunity, indicating a dual mechanism of action involving both direct tumor lysis and immune-mediated tumor suppression. NF27 is highly water-soluble at concentrations up to 40 mM. The peptide was rationally designed to exclude chemically reactive amino acids, enhancing its stability. Importantly, unlike many oncolytic peptides, NF27 exhibited minimal hemolytic activity, highlighting its safety profile. These attributes position NF27 as a promising candidate for the development of novel cancer therapeutics.

Compared to the three oncolytic peptides used in clinical trials, NF27 possesses unique characteristics. Unlike LTX-315 (*14*) and CyPep-1 (*15*), which contain nonnatural amino acids, NF27 is composed entirely of natural amino acids. This feature offers advantages such as lower synthesis costs and more predictable metabolic fates. Additionally, NF27 differs in peptide length and structural properties. Among existing oncolytic peptides, LTX-315 is the smallest, consisting of only 9 amino acids, while LL-37 (*13*) is the largest, with 37 amino acids. NF27 and CyPep-1 are similar in size, containing 23 and 27 amino acids, respectively. Structurally, LL-37 features a long α-helix with a short C-terminal tail, CyPep-1 forms a single α-helix, and LTX- 315 adopts a short helical structure, whereas NF27 contains a α-helix with a longer unstructured N-terminal region.

We demonstrated that NF27 significantly inhibits tumor growth and, in some cases, eradicates tumors completely. However, its efficacy varied across tumor models. NF27 completely eradicated CT26 tumors, whereas its effect on 4T1 tumors was relatively modest. While in vitro assays confirmed that CT26 cells were more sensitive to NF27 than other cell lines, the amount of NF27 injected into tumors exceeded the theoretical concentration required to induce cell death, suggesting that intrinsic cell sensitivity alone does not fully explain the observed differences in vivo. During intratumoral injection, we noticed that 4T1 tumors exhibited higher interstitial pressure than CT26 tumors, making injections more difficult in the former. Previous studies support this observation; Einen et al. reported that CT26 tumors contain less collagen than 4T1 tumors, leading to lower stiffness (*41*), while Muñoz et al. found that drugs administered intratumorally distribute more widely in softer tumors, such as B16, compared to stiffer tumors like MC38 (*42*). Based on these findings and our experience, we hypothesize that the discrepancy across tumor models may be influenced by intratumoral distribution of NF27.

In terms of clinical application, intratumoral injection of NF27 represents a promising strategy to reduce tumor burden while simultaneously activating anti-tumor immunity. Solubilized NF27 can be directly administered into transdermally accessible tumors, such as melanoma, breast cancer, and sarcoma. For tumors located within internal organs, image-guided techniques such as ultrasound-guided percutaneous injection (*43*) or endoscopic ultrasound (EUS)-guided fine needle injection (*44*) offer feasible routes for intratumoral delivery. Both approaches utilize real-time imaging to visualize internal anatomy and accurately guide needle placement for precise therapeutic injection.

Despite the promise of intratumoral delivery, several challenges remain. First, intratumorally administered drugs often exhibit rapid leakage from the injection site, limiting their retention and efficacy. Second, the significantly larger size and heterogeneity of human tumors compared to murine models pose difficulties to achieving uniform drug distribution throughout the tumor mass. Third, as a localized treatment, intratumoral injection is inherently limited in its capacity to address metastatic or disseminated disease.

To overcome these limitations, strategies to enhance intratumoral retention and distribution should be considered. These include incorporation of NF27 into hydrogel-based delivery systems (*45*), conjugation to tumor-binding ligands (*46*), or formulation within nanoscale drug carriers such as dendrimers, liposomes, or polymeric nanoparticles (*5*).

Additionally, given prior evidence of synergy between oncolytic peptides and immunotherapy, combining NF27 with immune checkpoint inhibitors may potentiate its immunostimulatory effects and improve therapeutic outcomes. Ultimately, the development of systemic delivery strategies for NF27 would address many of the constraints associated with intratumoral injection. For instance, gene delivery platforms (*47*) could be employed to enable tumor-selective expression of NF27, thereby extending its application to metastatic disease and expanding its translational potential.

## MATERIALS AND METHODS

### Study design

This study aimed to investigate the SAR of oncolytic peptides by evaluating the activity of CKS1-derived variants. We further assessed the therapeutic efficacy of NF27 in vitro and in vivo. In vitro assays were performed using murine and human cell lines, conducted in triplicate and repeated at least three times. Exceptions include evaluations of CKS1-derived peptides, where resource limitations prevented repeated experiments. Detailed experimental conditions are provided in the Materials and Methods section and in the corresponding figure legends. In vivo studies were conducted using BALB/c, C57BL/6, or BALB/c nude mice. Sample sizes were determined via power analysis in R. Randomization was performed prior to experimentation. For animal studies, mice with sufficient tumor size were randomized into treatment or control groups while blinded in a cloth-covered cage. Each group included at least five mice, and all experiments were independently repeated at least twice. In both in vitro and in vivo studies, no data or samples were excluded, and all data points are visualized in the figures.

### Peptides

All the peptides were synthesized by Genscript using a solid-phase peptide synthesis technique and provided as lyophilized powders. The C-terminus of all peptides was amidated if not described otherwise. For peptides used in the initial screening (NF04, NF05, NF06, NF07, NF08, NF09, NF10, NF11, NF12, NF13, NF14, NF15, NF16, NF17, NF18, NF19, NF20, NF21, NF22, NF23, NF24, NF25, NF26), the peptides were purchased as a purified peptide library. The purity was more than 80% as verified by HPLC and MS analyses. NF11 formed a hard gel upon exposure to water. All other peptides were solubilized in water. Peptides used in other assays (NF27, NF28, NF29, NF30, NF31, NF32, NF33, NF34, NF35, NF36, NF37, NF38, NF39, NF40, NF41, NF42, NF43) were synthesized individually and the purity was more than 90% as verified by HPLC and MS analyses. All these peptides were solubilized in water. For FITC-NF27, FITC was tagged on the N-terminus of NF27. The purity was more than 90% as verified by HPLC and MS analyses. Melittin was purchased from Genscript (RP20415).

### Cell Culture

4T1 murine mammary carcinoma cells (CRL-2539), CT26 murine colon carcinoma cells (CRL- 2638), B16-F10 murine melanoma cells (CRL-6475), NIH/3T3 normal murine fibroblast cells (CRL-1658), U2OS human osteosarcoma cells (HTB-96), HT-29 human colorectal cancer cells (HTB-38), and HepG2 human liver cancer cells (HB-8065) were purchased from the American Type Culture Collection. MCA205 murine fibrosarcoma cells (SCC173) were purchased from MilliporeSigma. HUVECs (CC-2519) and normal human lung fibroblasts (CC-2512) were purchased from LONZA. CT2A murine glioblastoma cells were a gift from Dr. Christopher Jackson. SUM149 cells were a gift from Dr. Zaver Bhujwalla. Panc10.05 human pancreatic cancer cells were a gift from Dr. Jackie Zimmerman. 4T1 cells, CT26 cells, and MCA205 cells were propagated in RPMI 1640 medium (Gibco, 11875119) supplemented with 10% FBS (MilliporeSigma, F4135). B16-F10 cells, CT2A cells, NIH/3T3 cells, U2OS cells, HT-29 cells, and HepG2 cells were propagated in DMEM (Gibco, 11965-092) supplemented with 10% FBS. SUM149 cells were propagated in DMEM/Nutrient Mixture F-12 Ham medium (Sigma-Aldrich, D6421) supplemented with 5 µg/mL human recombinant insulin (Gibco, 12585014) and 5% FBS. Panc10.05 cells were propagated in RPMI medium supplemented with 7 µg/mL human recombinant insulin and 10% FBS. HUVECs were propagated in EGM-2 BulletKit (LONZA, CC-3162). Normal human lung fibroblasts were propagated in FGM-2 BulletKit (LONZA, CC- 3132). All cell lines were incubated in 75cm^2^ Tissue Culture Flask (CELLTREAT Scientific Products, 229341) at 37°C in a 5% CO_2_ humidified atmosphere.

### LDH assay

1.0 × 10^4^ cells/well were seeded in a 96-well plate (Thermo Scientific, 15041) and incubated overnight. Serial dilution of peptides was prepared in the culture medium for each cell line. Cells were treated with the indicated concentration of peptides for 2 hours. CyQUANT LDH cytotoxicity assay (Invitrogen, C20301) was used following the manufacturer’s instructions to quantify the amount of LDH released in the culture medium. The LDH release was normalized to the amount of LDH released from the cells treated with lysis buffer. In experiments designed to generate dose–response curves, cells were treated with a range of concentrations, including doses low enough to elicit no detectable response and high enough to reach saturation of the treatment effect. To draw a dose-response curve, the LDH release was normalized to the LDH release of the highest concentration. The normalized LDH release was plotted on a log10 scale, and a dose-response curve was fitted using the L.4 function in the “drc” package in R. The “drc” package was used to calculate the EC_50_ values.

### Cell viability assay

1.0 × 10^4^ cells were seeded in a 96-well plate and incubated overnight. Serial dilution of peptides was prepared in the culture media for each cell line. Cells were treated with the indicated concentration of peptides for 2 hours. After peptide treatment, the cells were washed for 3 times with culture media. After the final wash, the media with exchanged with media containing 10% alamarBlue (Bio-Rad Laboratories, BUF012A), and the cells were incubated for 3 h. After a 1:5 dilution of the supernatant, the fluorescence (excitation/emission = 570 nm/610 nm) was measured using a plate reader (Agilent, BioTek Synergy H1). Finally, the cell viability was calculated by the following formula: cell viability (%) = 100 × (fluorescence from each sample – fluorescence from lysis buffer treated cells)/(fluorescence from non-treated cells – fluorescence from lysis buffer treated cells). In experiments designed to generate dose–response curves, cells were treated with a range of concentrations, including doses low enough to elicit no detectable response and high enough to reach saturation of the treatment effect. To draw a dose-response curve, the cell viability was normalized to the viability of the untreated cells. The normalized cell viability was plotted against concentration on a log10 scale, and a dose-response curve was fitted using the L.4 function in the “drc” package in R. The “drc” package was used to calculate the EC_50_ values.

### NCI-60 cancer cell line screening

NF27 was submitted to the National Cancer Institute Developmental Therapeutics Program and was evaluated for its anti-proliferative activity on a panel of 59 cancer cell lines. Out of the 60 cell lines, 1 cell line (SF-539) was excluded from the screening due to quality control issues. The five-dose screening was conducted using the NCI-60 classic protocol (https://dtp.cancer.gov/discovery_development/nci-60/methodology.htm) prior to the August 2023 methodology update. Briefly, each cell line was treated with 5 different concentrations of NF27 (0.01, 0.1, 1, 10, and 100 µM) for 48 hours. Cells were stained with Sulforhodamine B (SRB) after peptide treatment, and the number of viable cells was quantified. The cell growth was determined by comparing the number of viable cells before and after NF27 treatment. GI50 (drug concentration resulting in 50% reduction of cell growth), TGI (drug concentration resulting in total growth inhibition), and LC50 (drug concentration resulting in 50% reduction in the number of viable cells compared to before treatment), were reported based on the dose-response curve fitted against the experimental data.

### Acquisition of NCI-60 cell line molecular data

Z-score transformed mRNA expression data (exp) (*48*), protein expression data (swa) (*49*), and surface receptor expression data (sur) (*50*) were downloaded from the CellMinerCDB platform (*28*). The cell size data was acquired from Ortmayr et al, who estimated the sizes of 54 cell lines by analyzing their bright field images (*27*). The protein expression data and the surface receptor expression data were converted from the log2 intensity value to the raw intensity value and were subsequently transformed as Z-scores. The cell size data were also transformed as Z-scores.

### Correlation analysis of NF27 response data and NCI-60 cell line molecular data

Since the GI_50_ of melanoma cell line LOX IMVI was significantly higher compared to the other cell lines, we treated LOX IMVI as an outlier and removed the cell line from the correlation analysis. The GI_50_ values were converted to Z-scores. The Pearson correlation coefficient between the GI_50_ values (Z-score transformed) and the expression level of each molecule or the cell size (Z-score transformed) was calculated. Statistical testing was simultaneously conducted to determine the significance of the correlation. The p-values were corrected for multiple testing using the Benjamini-Hochberg procedure. We defined features that had a Benjamini-Hochberg adjusted p-value of less than 0.05 as features that significantly correlate with the GI_50_ values.

Gene enrichment analysis was conducted using the Database for Annotation, Visualization and Integrated Discovery (DAVID) platform (*51, 52*).

### Cell imaging

5.0 ×10^4^ cells were seeded in collagen-coated 35 mm glass-bottom dishes (MatTek, P35GCOL- 1.5-14-C) and incubated overnight. On the day of imaging, cells were stained with the appropriate dyes. Cytosolic staining was performed using Calcein-AM (Invitrogen, C1430) at a final concentration of 10 µM for 20 minutes. Nuclei were stained with Hoechst 33342 (Cell Signaling Technology, 4082) at a final concentration of 1 µg/mL for 10 minutes. Mitochondrial membrane potential was assessed by JC-1 staining (Invitrogen, T3168) at 5 µg/mL for 30 minutes. For each cell, the mitochondrial membrane potential was quantified by calculating the ratio of red (mitochondria with membrane potential) to green (mitochondria without membrane potential) fluorescence intensity. Mitochondria were stained with 1 µM MitoTracker Red FM (Invitrogen, M22425) for 30 minutes, and the ER was labeled with 1 µM ER-Tracker Red (Invitrogen, E34250) for 30 minutes. For Golgi apparatus staining, cells were incubated with 5 µM BODIPY FL C_5_-ceramide complexed to BSA (Invitrogen, B22650) at 4°C for 30 minutes, followed by a 30-minute incubation in fresh medium at 37°C. For experiments using FITC- NF27, cells were treated with 10 µM FITC-NF27 and imaged 30 minutes after treatment. Fluorescence imaging was performed using either a Zeiss LSM 780 or Zeiss LSM 880 with Airyscan confocal microscope. Brightfield imaging was performed using EVOS XL Core imaging system (Invitrogen, AMEX1000). Brightness and contrast of the images were adjusted appropriately using FIJI (*53*). Quantitative analysis of fluorescence intensity and localization was performed using CellProfiler (*54*).

### Caspase assay

1.0 × 10^4^ cells were seeded in 96-well plate and incubated overnight. Cells were treated with the indicated concentration of NF27 for 6 hours. Following treatment, cells were lysed using lysis buffer (Cell Signaling Technology, 7018). Caspase-3 activity in the lysates was measured using caspase-3 activity assay kit (Cell Signaling Technology, 5723) according to the manufacturer’s instructions. The signal intensity from untreated cells was subtracted from each condition to account for basal caspase-3 activity. Protein concentration in the lysates was determined using DC protein assay kit Ⅱ (Bio-Rad, 5000112) following the manufacturer’s instructions. Caspase-3 activity was normalized to total protein content and expressed as activity per µg of protein.

### Measurement of ATP release

4T1 cells or CT26 cells (1.0 × 10^4^ cells per wells) were seeded in 96-well plates and incubated overnight. The cells were treated with the indicated concentration of NF27 for 30 minutes. The supernatant was collected and the amount of ATP in the supernatant was quantified using the ENLITEN ATP assay system (Promega, FF2000), following the manufacturer’s instructions. ATP release was normalized to the amount of ATP released from the cells treated with lysis buffer.

### Measurement of cytokines from dendritic cells

4T1 and CT26 cells (2.0 × 10^5^ cells per well) were seeded in 6-well plates. DC2.4 cells (5.0 × 10^4^ cells per well) were seeded in 24-well plates. Cells were incubated overnight. 4T1 or CT26 cells were treated with 500 µL of 20 µM NF27 solution for 2 hours. After treatment, the conditioned media were collected, centrifuged, and filtered with Minisart syringe filter (Sartorius, 16553K). DC2.4 cells were treated with 250 µL of the conditioned media for 24 hours. The secretion levels of TNF-α and IL-6 were measured using ELISA kits specific for mouse TNF-α (antibodies.com, A719) and mouse IL-6 (antibodies.com, A78325), following the manufacturer’s instructions.

### Flow Cytometry

100 µL of 1.0 × 10^7^ cells/mL CT26 cells were subcutaneously injected in the flank of BALB/c mice. After 2 weeks, each tumor was intratumorally injected with 100 µL of 8 mg/mL NF27 (40 mg/kg body weight). 5 days after NF27 treatment, the tumors were excised, minced, and dissociated into single-cell suspensions using the gentleMACS Octo dissociator with heaters (Miltenyi Biotec, 130-134-029), gentleMACS C tubes (Miltenyi Biotec, 130-093-237), and tumor dissociation kit, mouse (Miltenyi Biotec, 130-096-730) in accordance with the manufacturer’s instructions. The cells were passed through a 70 µm cell strainer (Corning, 352350). The red blood cells were lysed using RBC lysis buffer (10X) (BioLegend, 420301).

Cells were then stained using the LIVE/DEAD fixable aqua dead cell stain kit (Invitrogen, L34957). Surface staining was performed using the antibodies in **Table S1**. Fluorescence intensities were acquired using the Cytek Aurora system (Cytek Biosciences) and subsequently analyzed to quantify each immune cell population.

### Animal models

The protocols used in this study were approved by the Institutional Care and Use Committee at Johns Hopkins Medical Institutions. Four- to eight-week-old female BALB/c mice were obtained from Charles River (028). For the HT-29 model, four- to eight-week-old BALB/c nude mice were obtained from Charles River (194). The cancer cells were harvested and suspended in PBS, and 100 µL of the cell suspension was injected into mice. For the CT26 model, 1.0 × 10^6^ CT26 cells were injected subcutaneously in the left flank. For the 4T1 model, 2.5 × 10^4^ 4T1 cells were injected into the first mammary fat pad. For the B16-F10 model, 1.0 × 10^5^ cells were injected subcutaneously into the left flank. For the HT-29 model, 1.0 × 10^6^ HT-29 cells were injected into the left flank. After 1 week, the animals were randomized into control and NF27 treatment groups. We intratumorally injected 50 µL of 16 mg/mL NF27 (40 mg/kg body weight) every 2 days. For the control group, an identical volume of water was injected intratumorally. The tumor size was measured by using a caliper, and the volume was calculated by using the formula 0.52 × (length) × (width).

### Mass spectrometry

The pharmacokinetics of NF27 in murine serum was measured by Origin Bioanalytical laboratory. Briefly, NF27 was incubated in murine serum containing K_2_EDTA as anticoagulant at the indicated time and condition. NF27 and its respective internal standard NF31 were extracted from 20 µL of serum using protein precipitation extraction. The extracts were evaporated to dryness, reconstituted, and then analyzed by LC-MS/MS.

### Hemolysis assay

Packed human red blood cells (BioChemed) were transferred to tubes containing sterile PBS and washed by centrifugation at 1700g for 10 min until the supernatant was clear. Pelleted RBCs were then resuspended in PBS to create a 2% RBC suspension. Peptides were prepared at twice the final concentration (2× peptide solution) by serial dilution in PBS. In a V-bottom 96-well plate, 50 μL of the 2% RBC suspension was combined with 50 μL of 2× peptide solution. 1% Triton-X in PBS (final concentration) was used as the positive control. The 96-well plate was incubated for 1 h at 37°C in a 5% CO₂ cell culture incubator. After incubation, the entire plate was spun at 1700g for 5 min. The supernatants were transferred to a clean flat-bottom 96-well plate and the absorbance was read at 405 nm. Percentage hemolysis was calculated as (A_sample_ – A_PBS_)/(A_TritonX_ – A_PBS_)·100.

### Prediction, visualization, and the analysis of peptide structure

Peptide structures were predicted using AlphaFold2 via the ColabFold platform with default parameters (*55*). Electrostatic surface potentials were calculated using the APBS plugin in PyMOL. The predicted structures, along with their electrostatic properties and hydrophobicity, were visualized in PyMOL. Physicochemical properties of the peptides were computed using the “modlAMP” package in Python (*24*). The helical wheel projections of the peptides were visualized using a Python script developed by Saladi (*56*).

### Statistical analysis

All statistical analyses were performed using R. Differences between two groups were assessed using unpaired two-tailed Welch’s t-test, with p-values < 0.05 considered statistically significant. For comparisons between the control and multiple treatment groups, Dunnett’s test was applied. Two-way ANOVA was used to evaluate the effect of treatment in *in vivo* tumor growth studies. If not stated otherwise, all data represent at least three independent experiments, and results are expressed as mean ± standard error of the mean (SEM). Individual data points from each experiment were plotted. In cases with significant variability across experiments, representative data from a single experiment are shown, with technical replicates plotted where applicable.

## Acknowledgments

The authors thank the National Cancer Institute Developmental Therapeutics Program (NCI/DTP) https://dtp.cancer.gov for providing screening data of NF27 present in this manuscript (NSC# 848831). Fluorescent microscopy was performed at the Microscope Facility of Johns Hopkins University, which receives financial support from NIH grant S10OD016374 and S10OD023548. Flow cytometry was performed by Ross Flow Cytometry Core at Johns Hopkins University which receives financial support from NIH grant S10OD026859.

## Author contributions

Conceptualization: NF; Funding acquisition: NF, ACM, NBP, ASP; Investigation: NF, ARC, WY, AP; Methodology: NF, ACM, NBP, ASP; Project administration: ASP; Supervision: ESC, ACM, NBP, ASP; Visualization: NF; Writing – original draft: NF; Writing – review & editing: NF, ESC, ACM, NBP, ASP

## Data and materials availability

All data are available in the main text or the supplementary materials.

