## Supplementary Materials for "Oncolytic peptide NF27 effectively inhibits tumor growth and eradicates tumors in multiple cancer types"

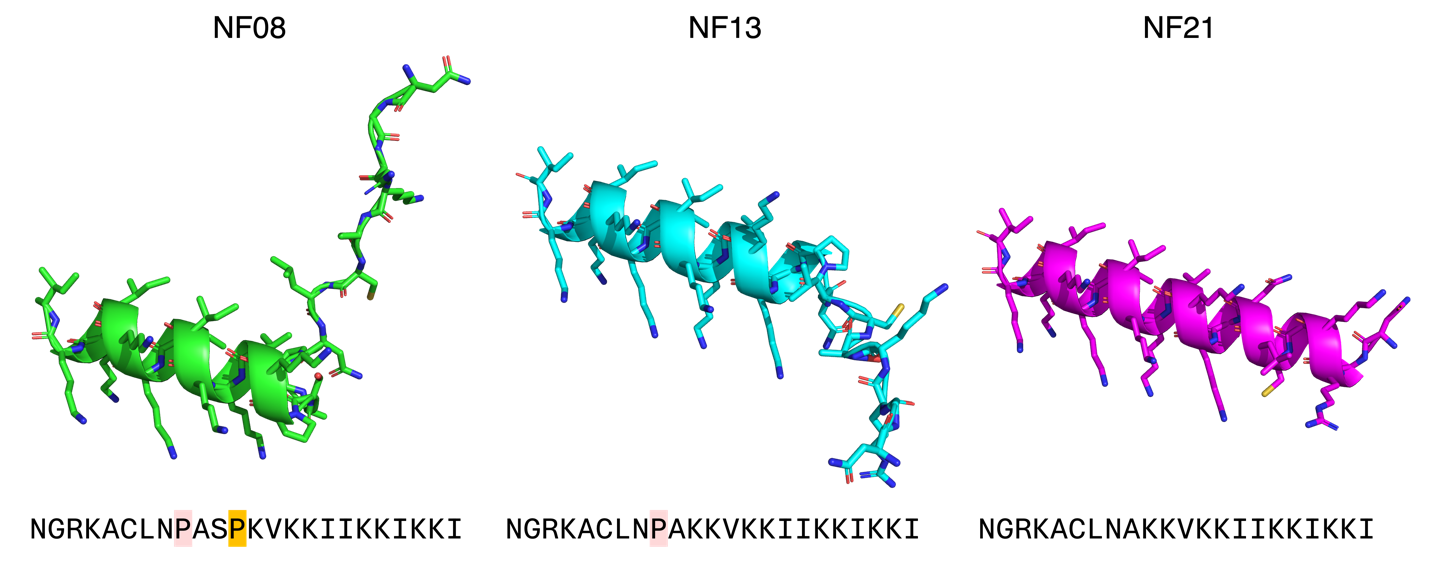

**Fig. S1. Structures of NF08, NF13, and NF21.** The structures of the three peptides were predicted using AlphaFold2. The differences between the peptides consist of the highlighted proline residues.

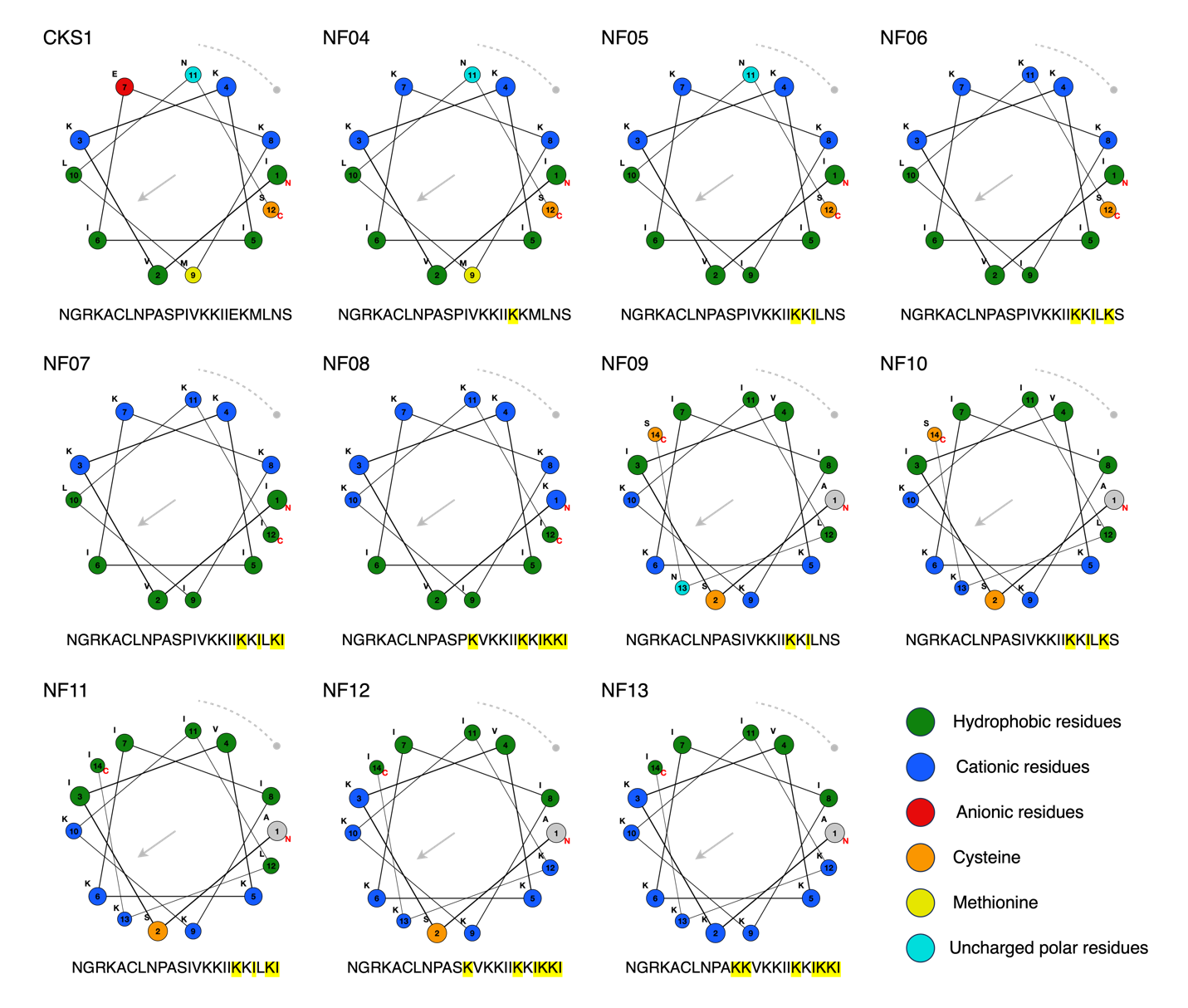

**Fig. S2. Helical wheel projections of the CKS1-derived peptides.** For CKS1 – NF08, the helical region after P12 is visualized. For NF09 – NF13, the helical region after P9 is depicted. The amino acids that differ from the original CKS1 sequence are highlighted in yellow.

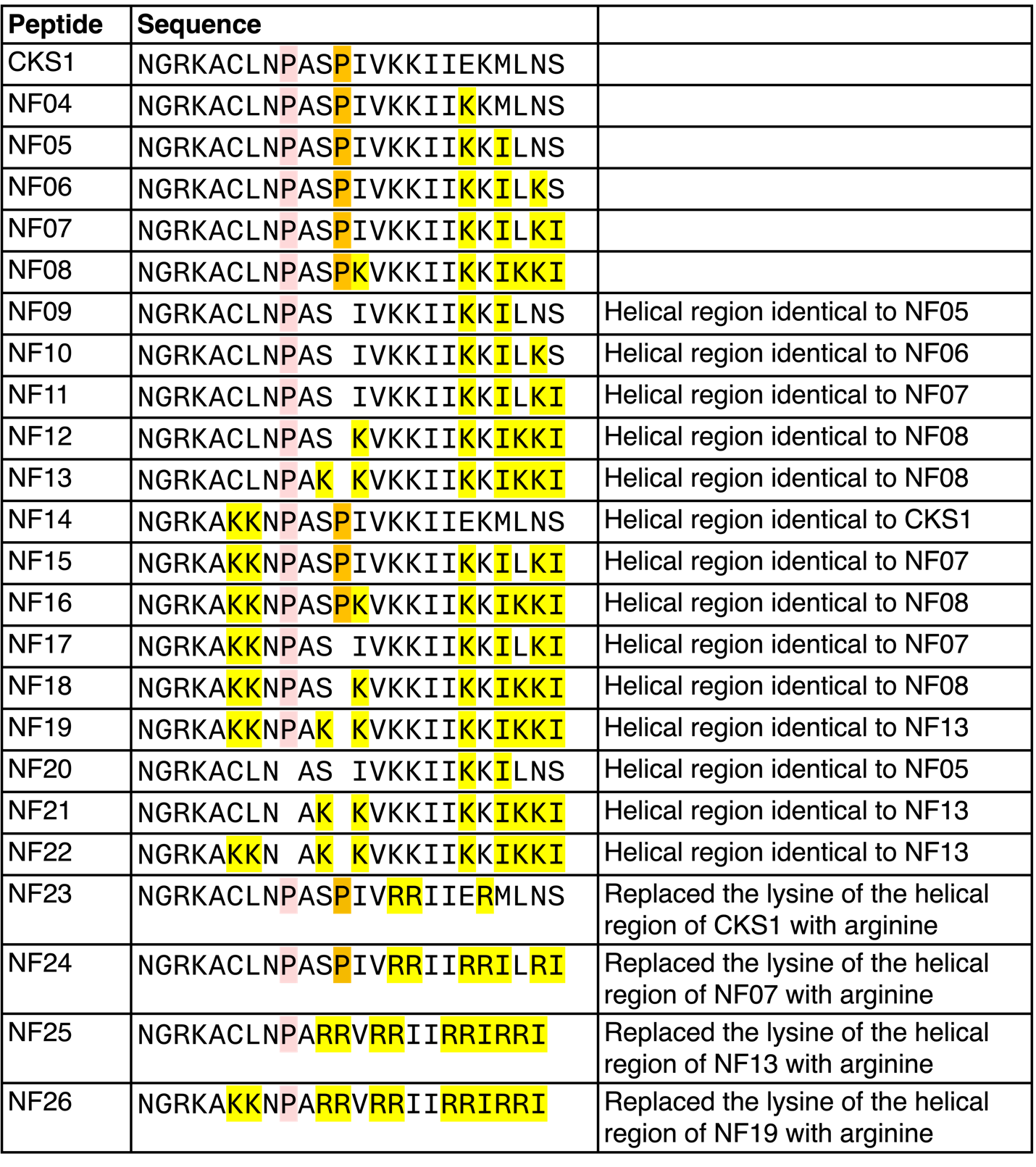

**Fig. S3. Sequence alignment of CKS1-derived peptides.** The two proline residues are highlighted in pink and orange. The amino acids that differ from the original CKS1 sequence are highlighted in yellow.

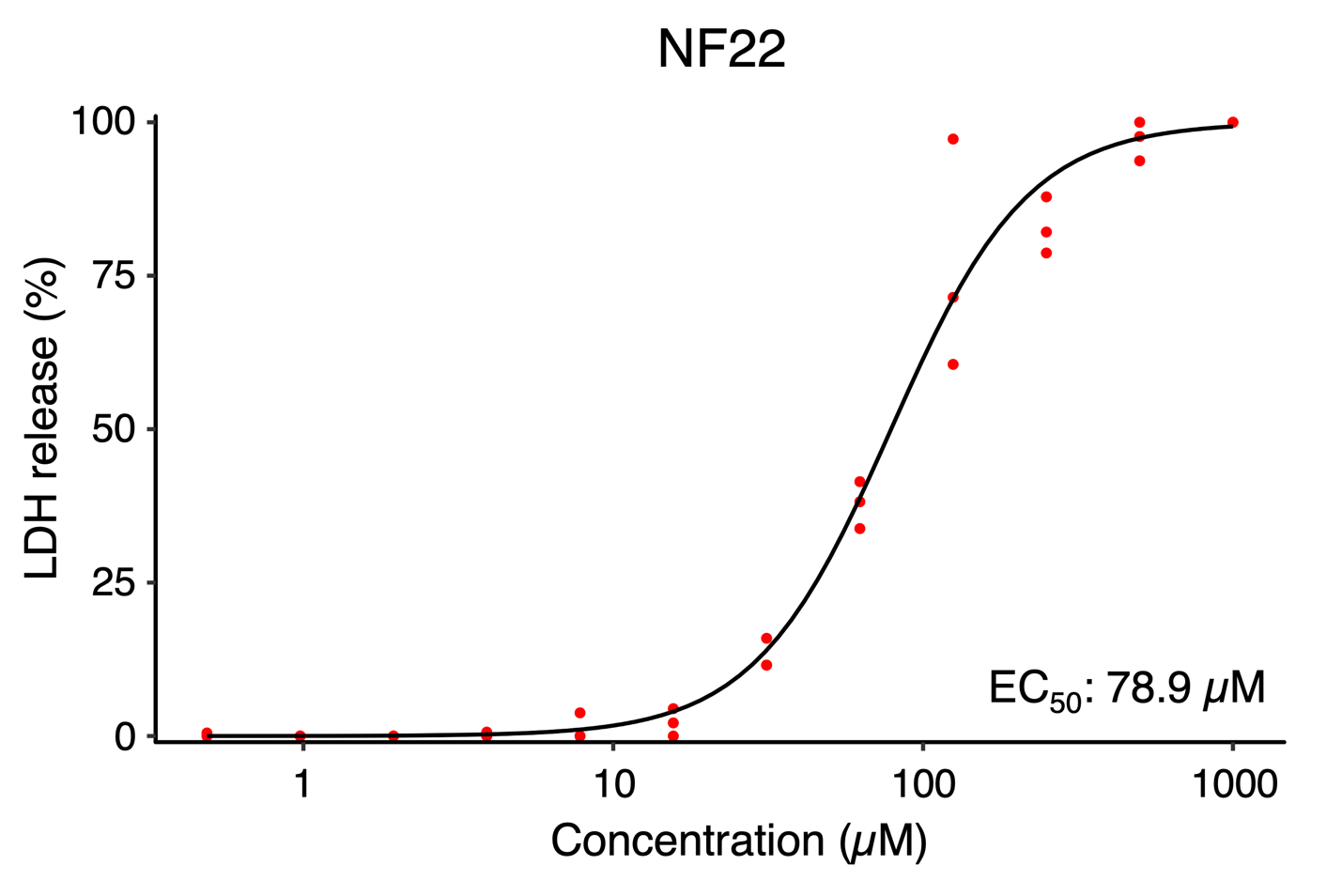

**Fig. S4. The oncolytic activity of NF22.** The release of LDH from 4T1 cells treated with NF22. Representative of N = 3. Each data point represents a technical replicate.

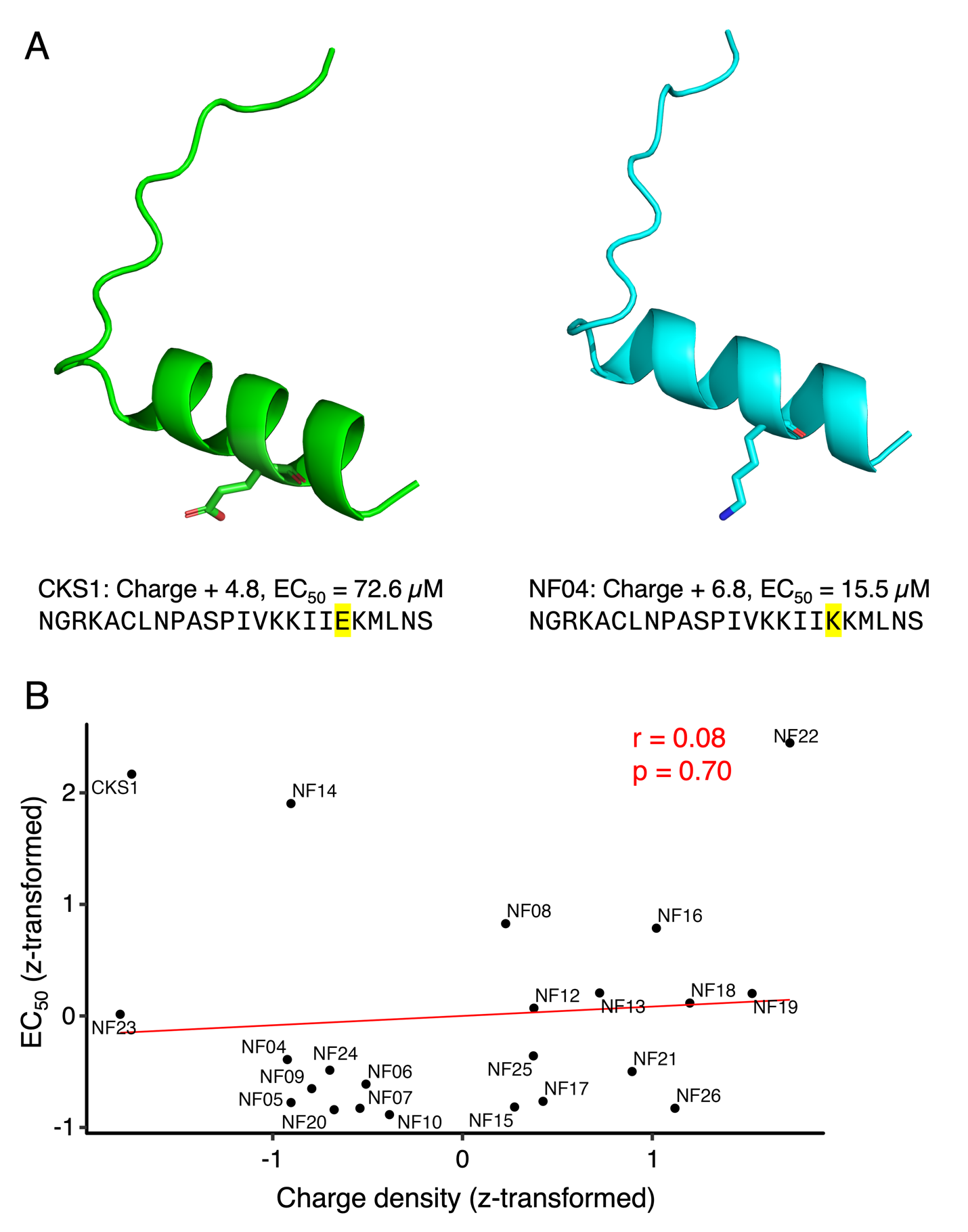

**Fig. S5. The impact of positive charge on oncolytic activity.** (A) The predicted structures and the properties of CKS1 and NF04. Replacing a glutamic acid residue with lysine residue enhanced the oncolytic activity. (B) The EC_50_ values were plotted against the charged density. The Pearson correlation coefficient (r) and the p-value were calculated to assess the significance of the correlation.

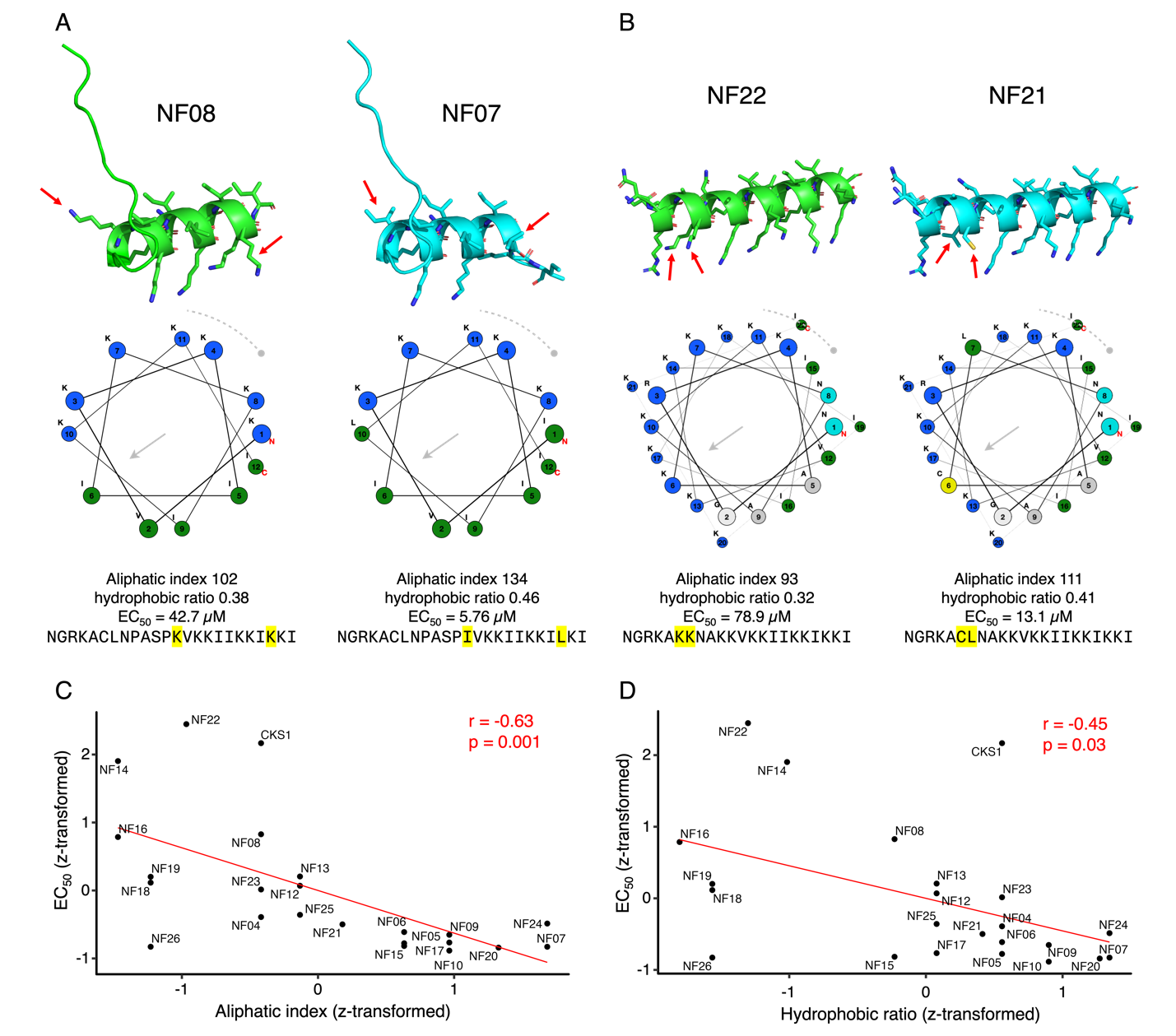

**Fig. S6. The impact of hydrophobicity on oncolytic activity.** (A) The predicted structures and the properties of NF08 and NF07. Sequential differences between the two peptides are highlighted in yellow, with red arrows indicating their locations in the predicted structures. NF07 features a larger hydrophobic region and exhibits higher oncolytic activity than NF08. (B) The predicted structures and the properties of NF22 and NF21. (C) The correlation between the EC_50_ values and the aliphatic index of each peptide. The Pearson correlation coefficient (r) and the p-value were calculated to assess the significance of the correlation. (D) The correlation between the EC_50_ values and the hydrophobic ratio of each peptide.

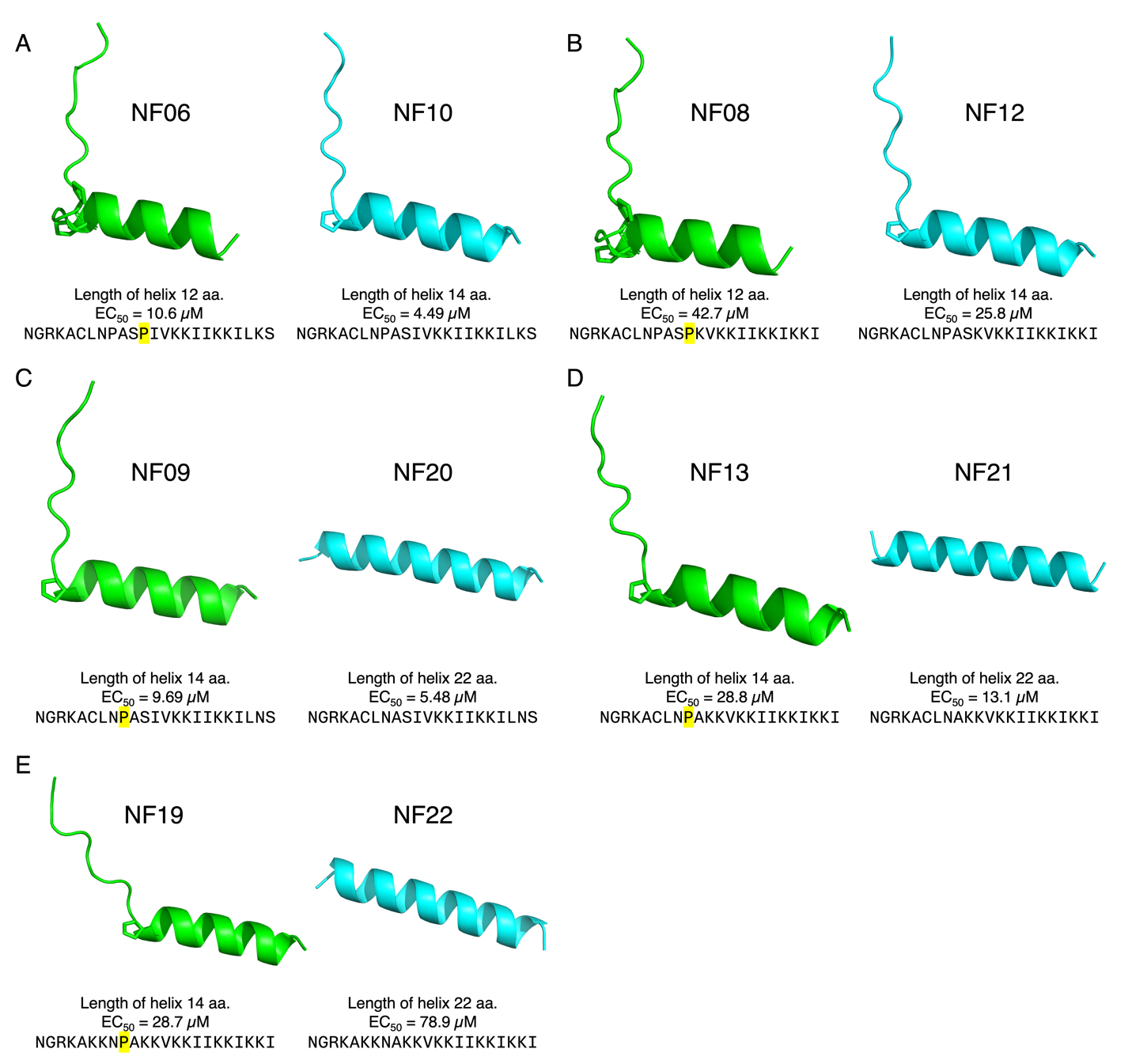

**Fig. S7. The impact of proline residues and the length of the helical region on oncolytic activity.** (A – D) The predicted structures and the properties of each peptide. Peptides on the right, which have longer helical regions due to the removal of proline residues, exhibit lower EC_50_ values compared to those on the left. (E) The predicted structures and the properties of NF19 and NF22. In this case, NF22 has a higher EC_50_ value compared to NF19.

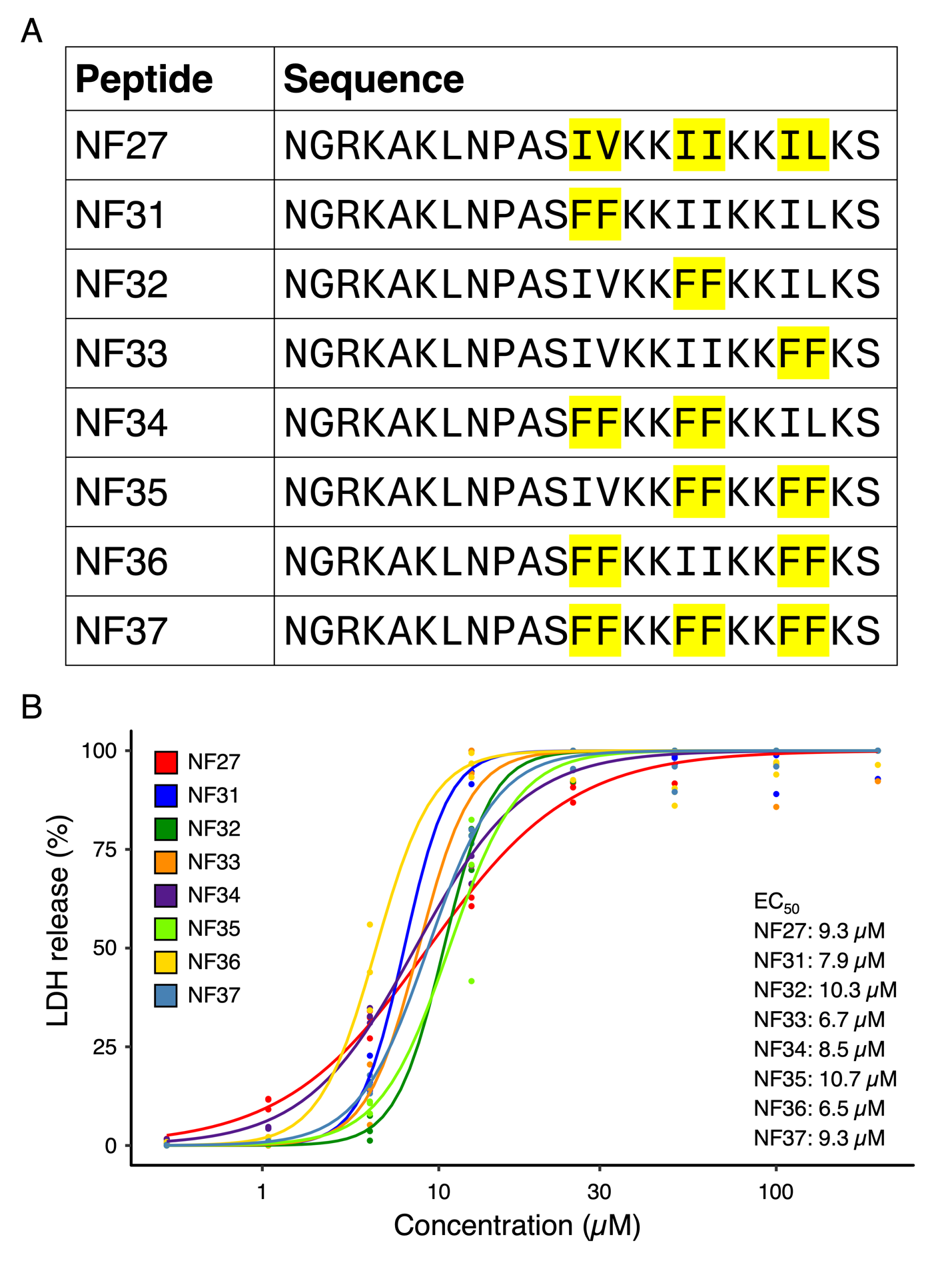

**Fig. S8. Activity of peptides using phenylalanine residues.** (A) The sequential alignment of NF27 and NF31 – NF37. The amino acids that differ from NF27 are highlighted in yellow. (B) Dose-response curves and EC_50_ values of NF27. 4T1 cells were treated with each peptide, and cytotoxicity was assessed by quantifying LDH release.

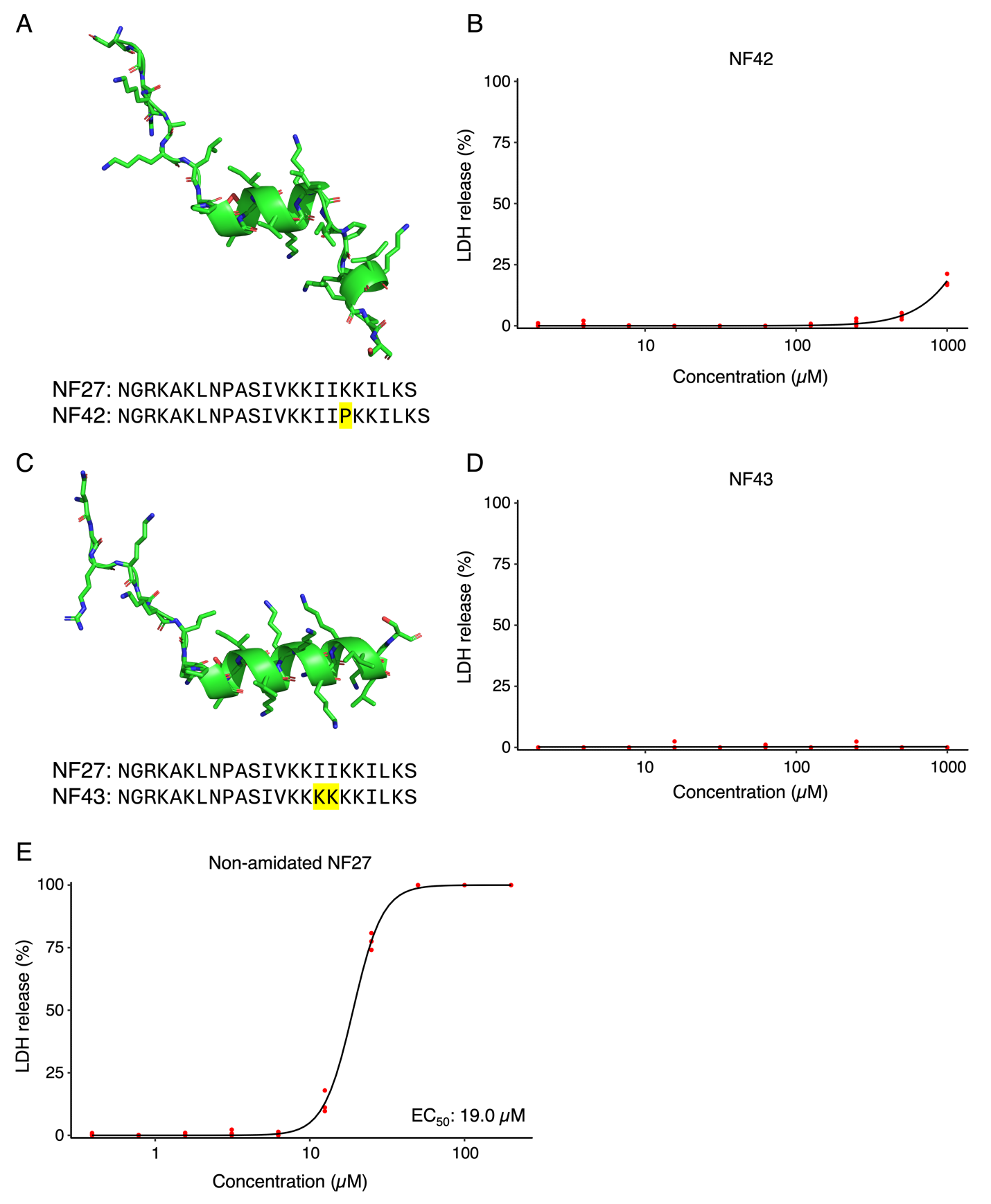

**Fig. S9. Structures and activities of NF42, NF43, and non-amidated NF27.** (A) The predicted structure of NF42. (B) The LDH release from 4T1 cells treated with NF42. (C) The predicted structure of NF43. (D) The LDH release from 4T1 cells treated with NF43. (E) The LDH release from 4T1 cells treated with non-amidated NF27.

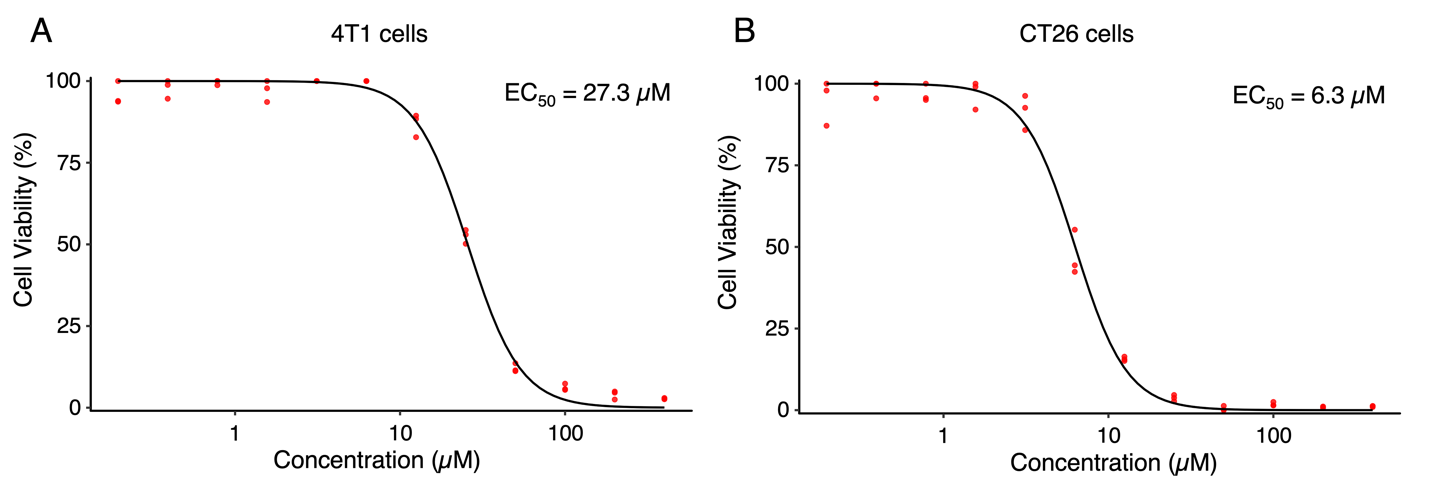

**Fig. S10. Oncolytic activity of NF27 confirmed by cell viability assay.** (A) Cell viability of 4T1 cells treated with NF27, assessed using an alamarBlue assay. Cell viability was normalized to that of 4T1 cells treated with the lowest NF27 concentration. A dose-response curve was fitted to experimental data to determine the EC_50_ value. (B) The cell viability of CT26 cells treated with NF27.

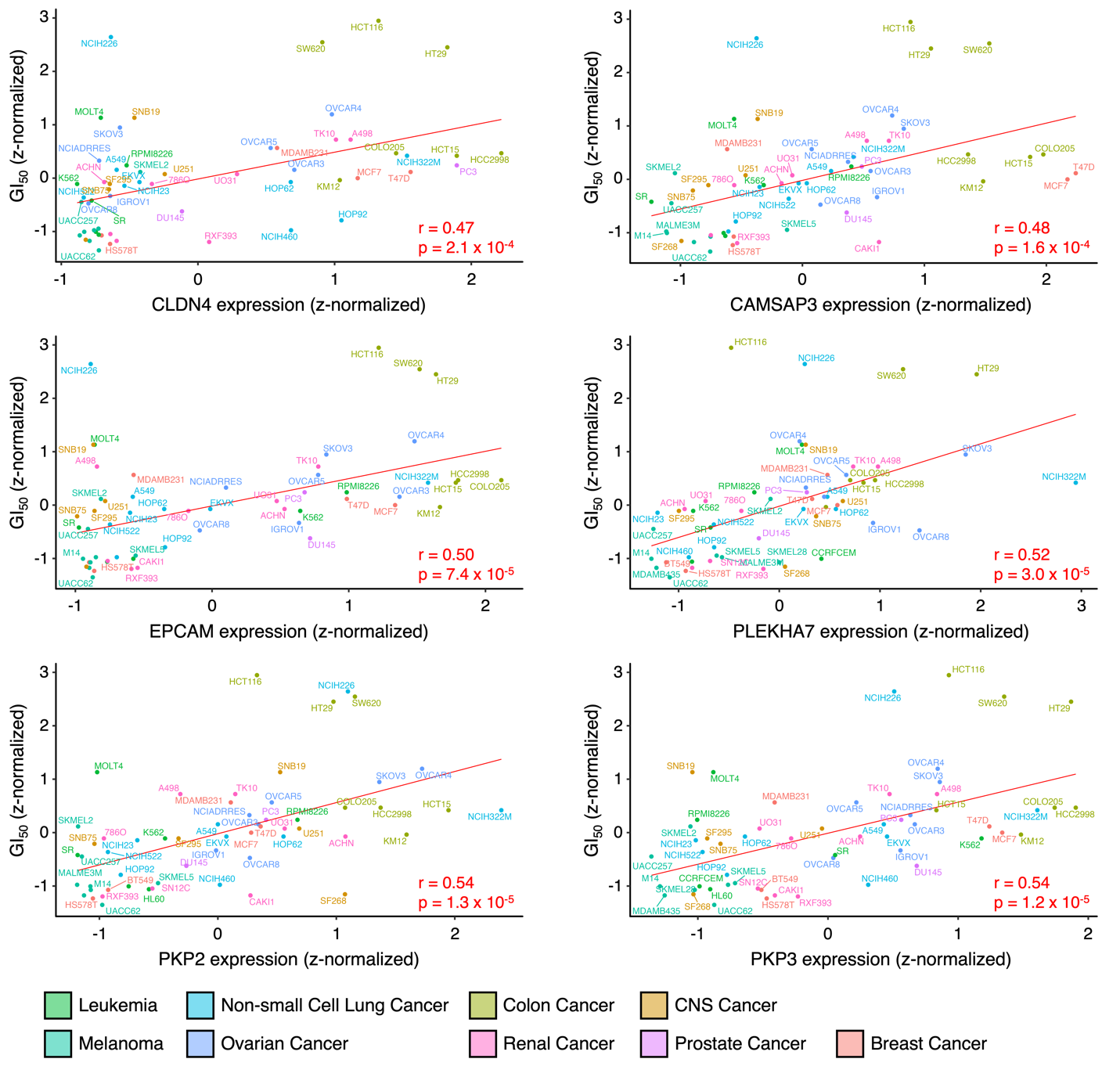

**Fig. S11. Correlation between the expression of cell-cell adhesion-related genes and the GI_50_ values.** Scatter plots of z-normalized expression levels of claudin 4 (CLDN4), calmodulin regulated spectrin associated protein family member 3 (CAMSAP3), epithelial cell adhesion molecule (EPCAM), pleckstrin homology domain containing A7 (PLEKHA7), plakophilin 2 (PKP2), and plakophilin 3 (PKP3) against z-normalized GI_50_ values. The Pearson correlation coefficient (r) and corresponding p-values were calculated to assess the significance of the correlation.

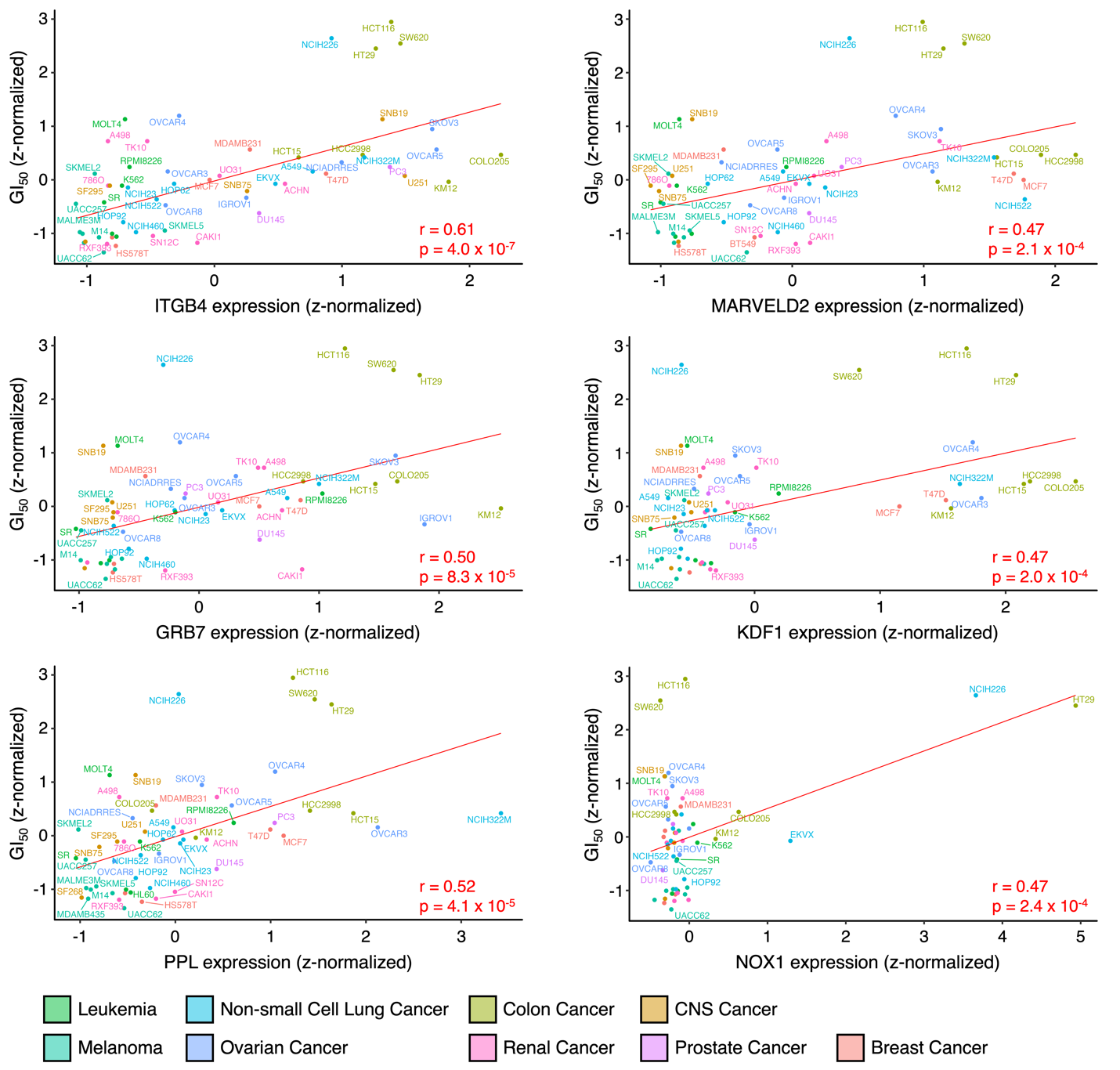

**Fig. S12. Correlation between the expression of cell-cell adhesion-related genes and the GI_50_ values (continued).** Scatter plots of z-normalized expression levels of integrin subunit beta 4 (ITGB4), MARVEL domain containing 2 (MARVELD2), growth factor receptor-bound protein 7 (GRB7), keratinocyte differentiation factor 1 (KDF1), periplakin (PPL), and NADPH oxidase 1 (NOX1) against z-normalized GI_50_ values. The Pearson correlation coefficient (r) and corresponding p-values were calculated to assess the significance of the correlation.

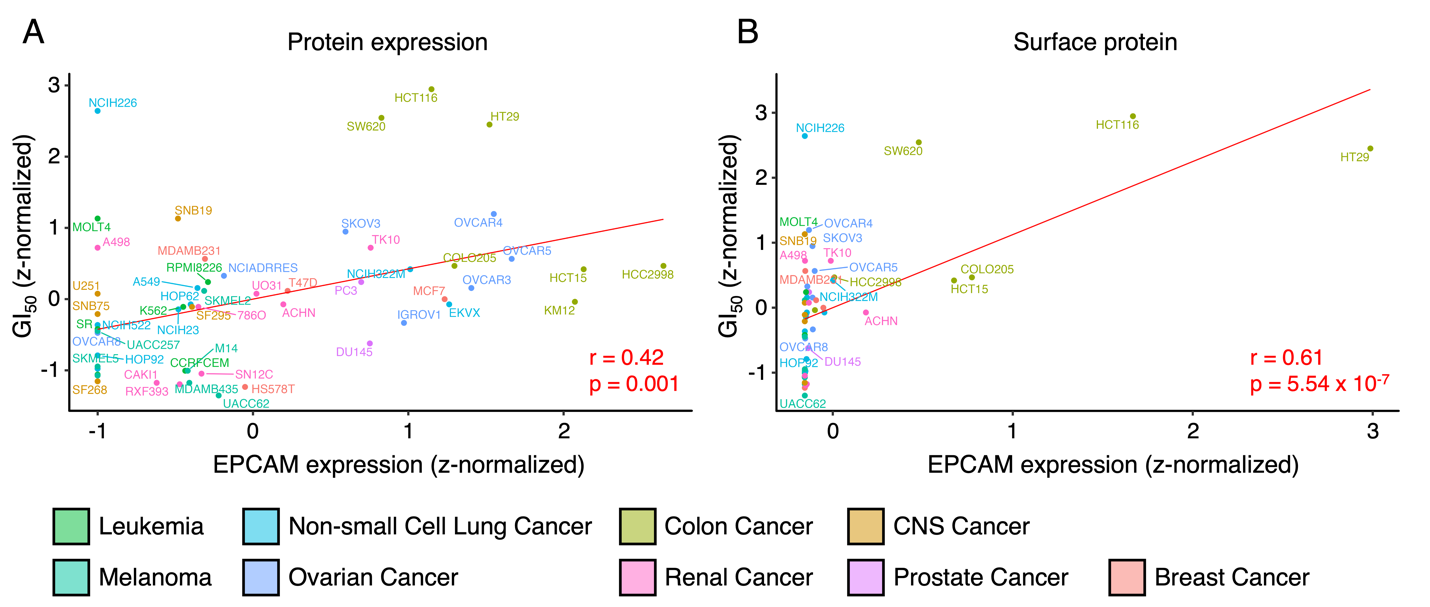

**Fig. S13.** **Correlation between the expression of EPCAM protein and the GI_50_ values.** (A) Scatter plot showing the correlation between z-normalized EPCAM protein expression and z-normalized GI_50_ values. The Pearson correlation coefficient (r) and corresponding p-values were calculated to assess the significance of the correlation. (B) Scatter plot showing the correlation between z-normalized EPCAM protein expression on the cell surface and z-normalized GI_50_ values.

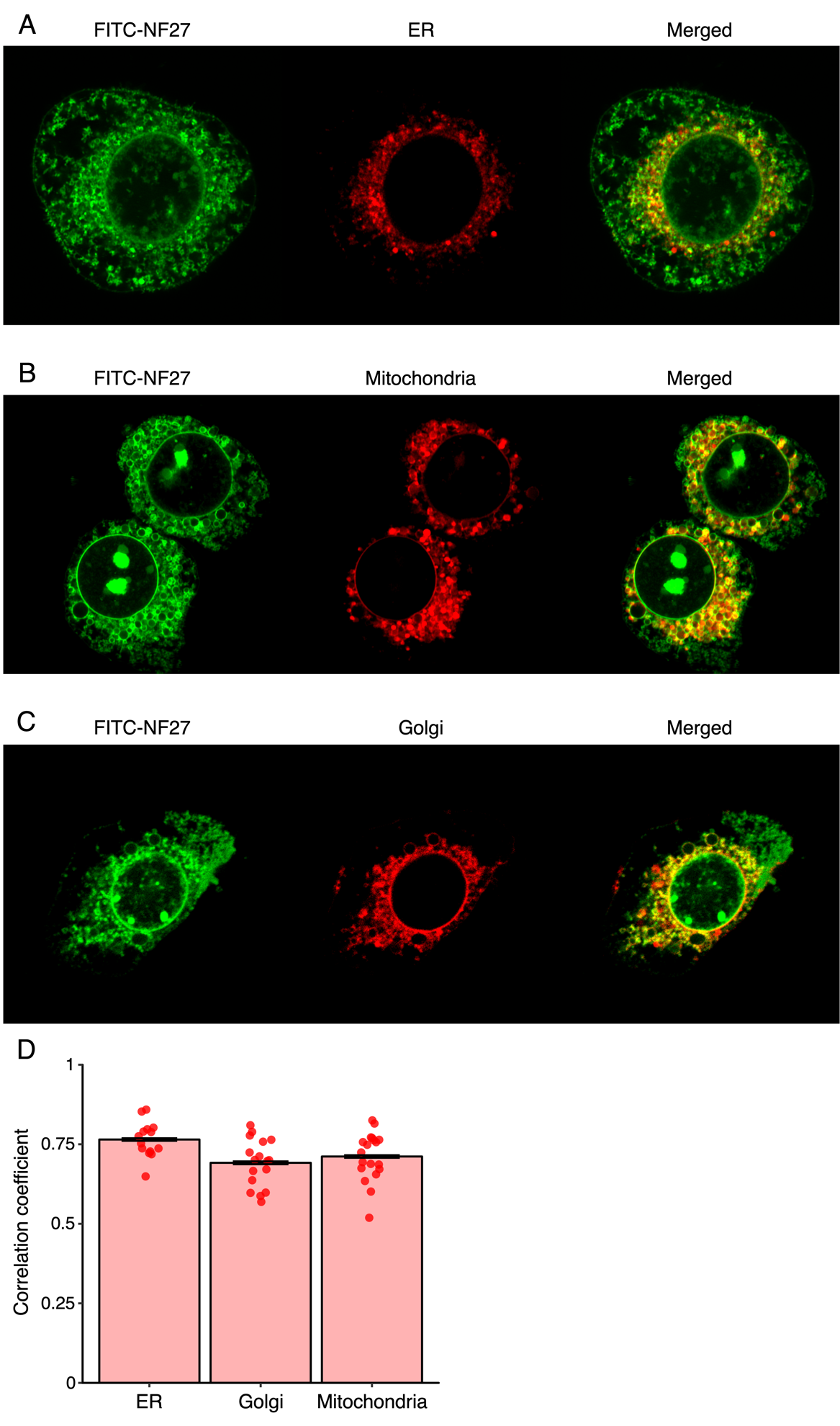

**Fig. S14. Co-localization of FITC-NF27 with organelles.** Representative fluorescence images showing the co-localization of FITC-NF27 with (A) ER, (B) Golgi apparatus, and (C) mitochondria. (D) Quantification of co-localization using Pearson correlation coefficients between FITC and organelle-specific markers. Each data point represents the coefficient calculated from an individual cell.

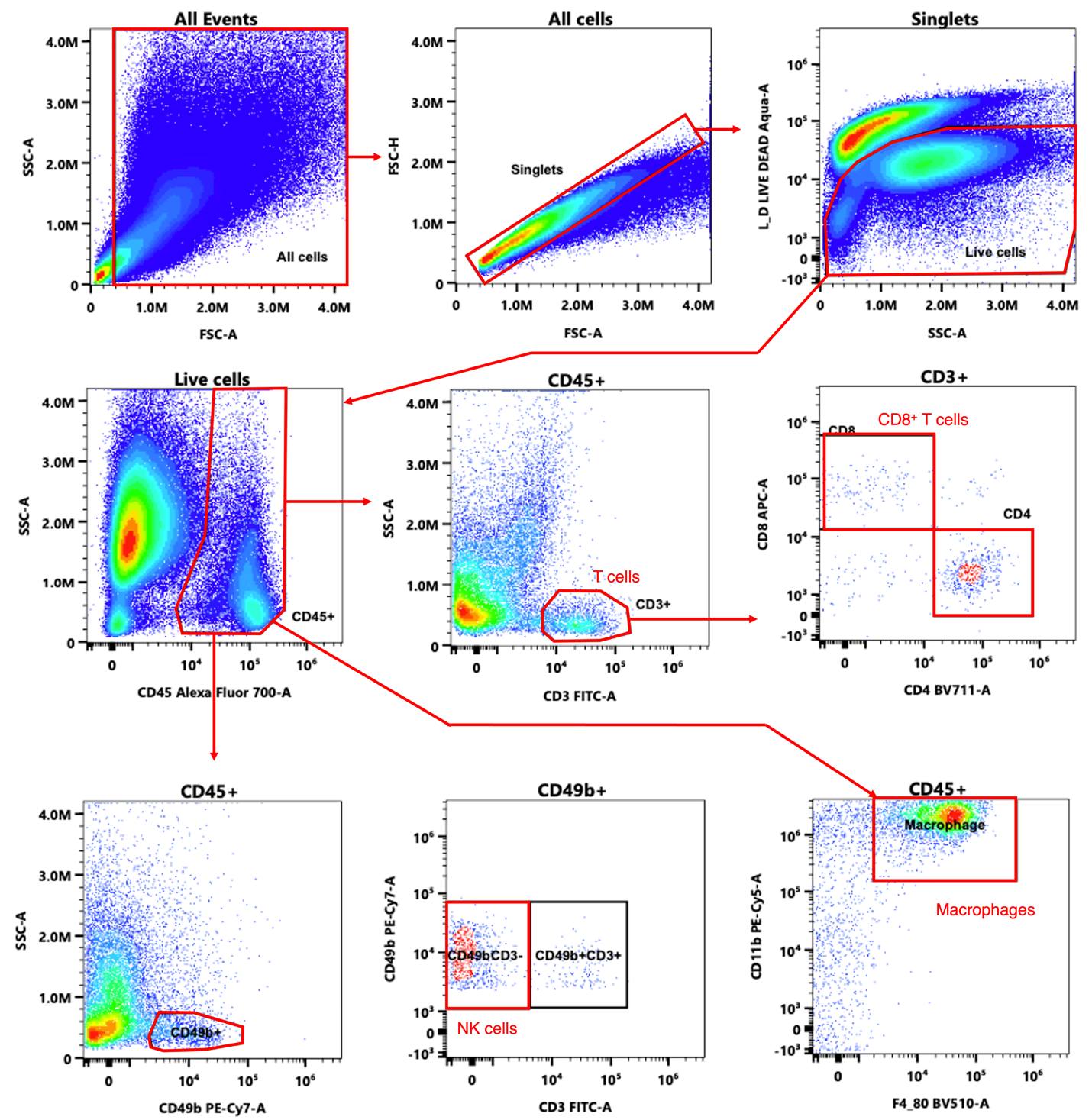

**Fig. S15. Gating strategy for the flow cytometry experiments.** T cells were defined as CD45^+^CD3^+^ cells, NK cells were defined as CD45^+^CD49b^+^CD3^-^ cells, macrophages were defined as CD45^+^CD11b^+^F4/80^+^ cells.

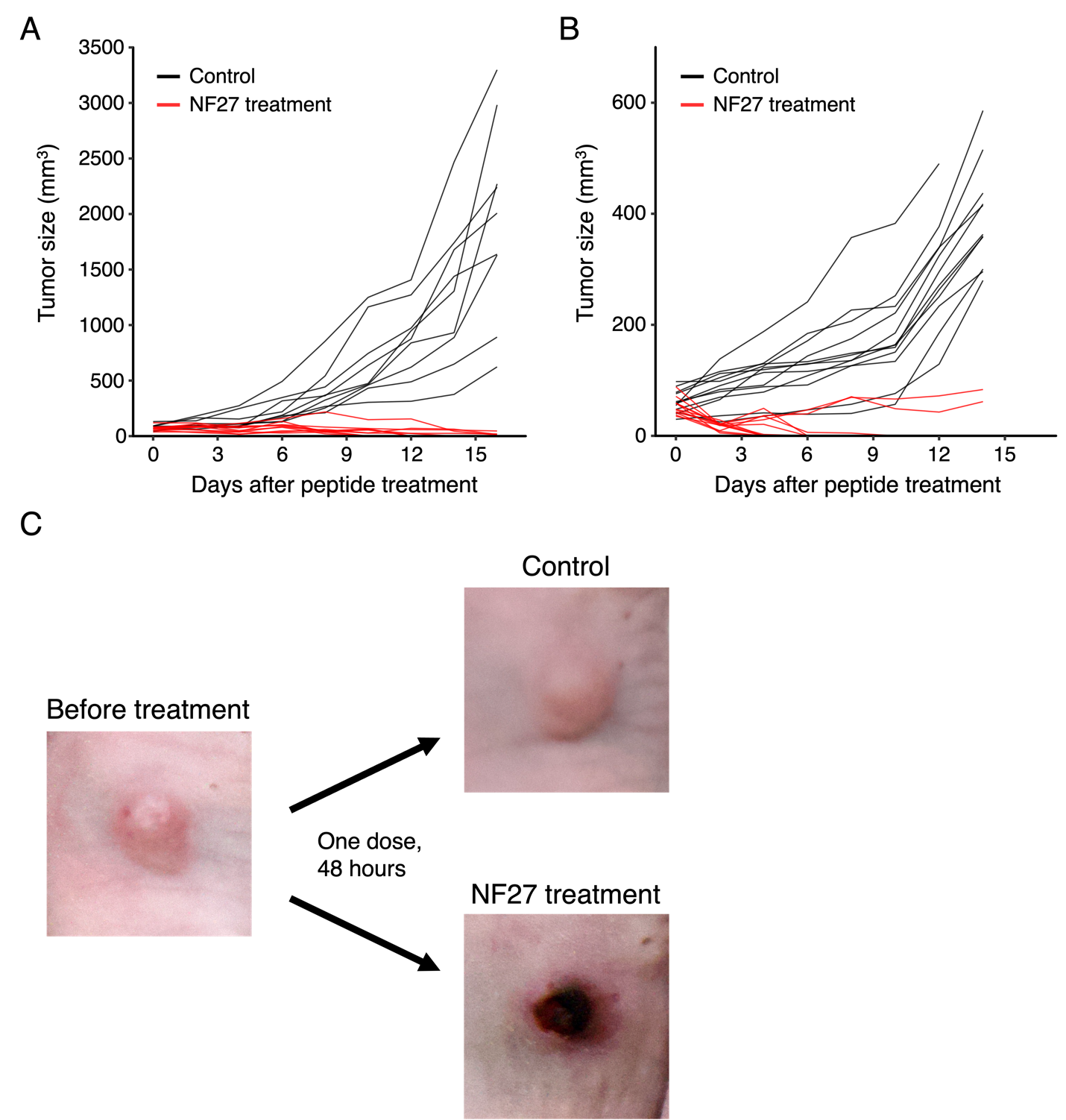

**Fig. S16.** **NF27 eradicates tumor.** (A) Spider plot showing individual CT26 tumor growth trajectories. (B) Spider plot showing individual HT-29 tumor growth trajectories. (C) Representative images of HT-29 tumor following a single intratumoral injection of vehicle or NF27.

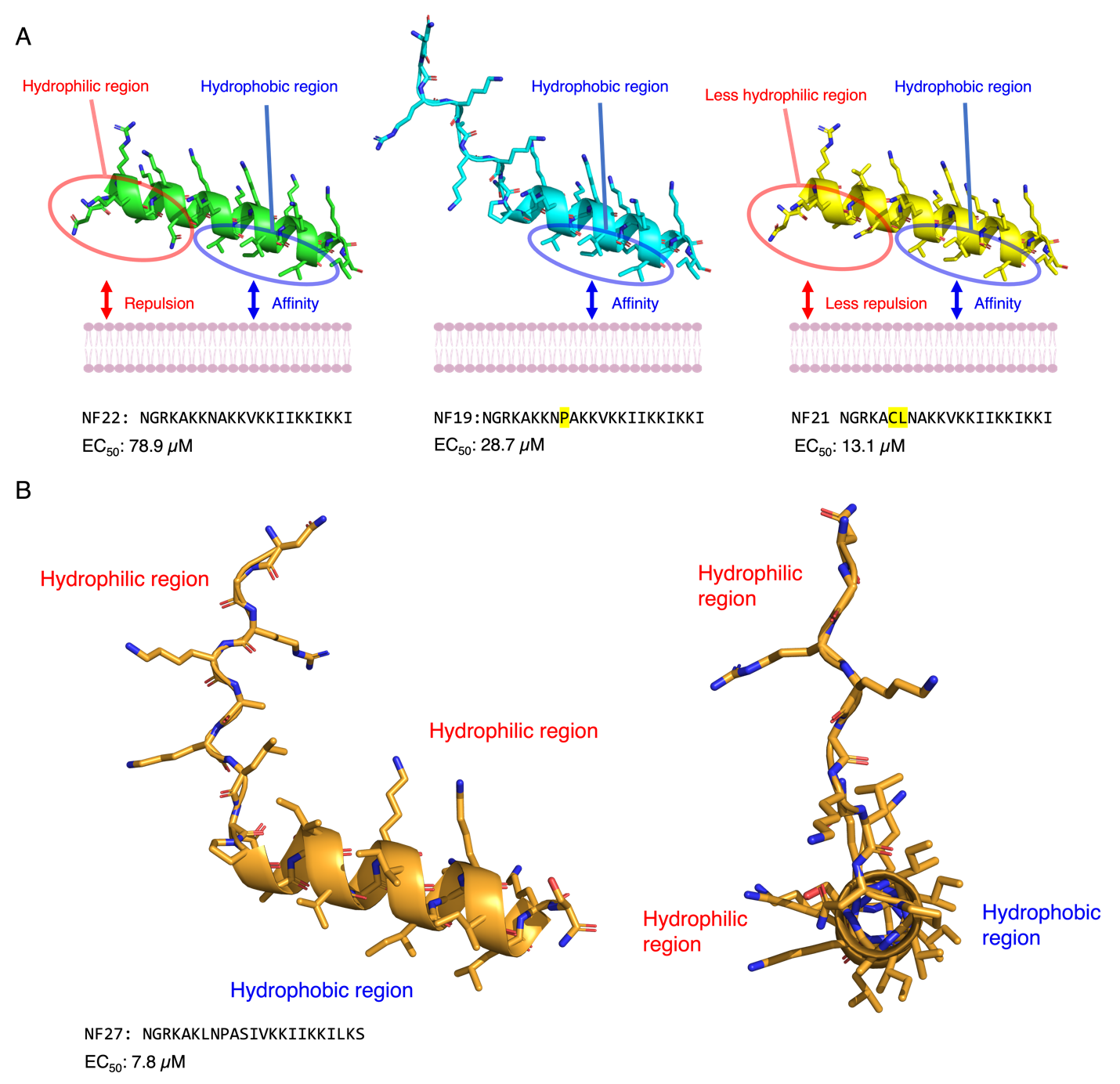

**Fig. S17. Hypothetical model explaining the differences between NF19, NF21, and NF22.** (A) Illustration of the spatial relationship between hydrophilic and hydrophobic regions in each peptide. In NF22, the hydrophilic and hydrophobic regions align on the same side of the helix, potentially increasing repulsive interactions with the plasma membrane. In contrast, the proline-induced kink in NF19 separates these regions, reducing repulsion. Due to substitutions of hydrophilic amino acids to hydrophobic amino acids, NF21 has less hydrophilicity in the hydrophilic region and can potentially enhance membrane interaction through reduced repulsion and increased hydrophobic interaction. (B) The structure of NF27 illustrating the location of the hydrophilic and the hydrophobic regions.

| **Antigen** | **Fluorophore** | **Company** | **Clone** | **Catalog Number** |
| --- | --- | --- | --- | --- |
|  | LIVE/DEAD Aqua | Invitrogen |  | L34957 |
| F4/80 | BV510 | BioLegend | BM8 | 123135 |
| CD4 | BV711 | BioLegend | GK1.5 | 100447 |
| CD3ε | FITC | BioLegend | 145-2C11 | 100305 |
| CD11b | PE/Cy5 | BioLegend | M1/70 | 101209 |
| CD49b | PE/Cy7 | BioLegend | DX5 | 108921 |
| CD8 | APC | BioLegend | 53-6.7 | 100711 |
| CD45 | Alexa Fluor 700 | BioLegend | 30-F11 | 103127 |

**Table S1. Antibodies used in this study.**
